# Growth capacity and mortality burden shape microbial persistence under osmotic stress in *Bacteroides thetaiotaomicron*

**DOI:** 10.64898/2026.09.08.750096

**Authors:** Hans Ghezzi, Ricky Wolff, Asmita Jain, Juan C. Burckhardt, Michael Hunter, Katharine M. Ng, Shaqed Carasso, Naama Geva-Zatorsky, Nandita Garud, Carolina Tropini

**Affiliations:** Department of Bioinformatics, University of British Columbia, Vancouver, Canada; Department of Ecology and Evolutionary Biology, University of California, Los Angeles, USA; Department of Microbiology & Immunology, University of British Columbia, Vancouver, Canada; Department of Cell Biology and Cancer Science, Rappaport Faculty of Medicine and Research Institute, Rappaport Technion Integrated Cancer Center (RTICC), Technion-Israel Institute of Technology, Haifa, Israel; Humans and the Microbiome Program, Canadian Institute for Advanced Research (CIFAR), Toronto, Canada; School of Biomedical Engineering, University of British Columbia, Vancouver, Canada

**Keywords:** Microbiome, osmolality, growth rate, mortality, adaptation, *Bacteroides thetaiotaomicron*, phase variation, persistence

## Abstract

Microbial persistence in the gut is central to sustaining beneficial host-microbiota interactions; however, characterizing determinants of persistence *in vivo* can be challenging due to the complexity of native microbial communities interacting with the host. Here, we used a defined microbial consortium to demonstrate that intrinsic *in vitro* growth capacity becomes increasingly predictive of *in vivo* relative abundance as intestinal osmolality increases. In *Bacteroides thetaiotaomicron*, *in vivo* abundance was higher than predicted from growth capacity alone, indicating that additional physiological factors contribute to persistence. We identify mortality as one such determinant: adaptation to prolonged osmotic stress markedly reduced cell loss without enhancing growth capacity. This adaptive phenotype was associated with rapid and reversible phase variation in cell surface-associated PUL78/80 rather than fixed *de novo* mutations, with the PUL78/80-OFF state becoming progressively enriched among surviving cells as mortality increased. Our findings demonstrate that in addition to growth capacity, reduced cell loss can provide a distinct route to microbial persistence during environmental perturbation.

## Introduction

The human gastrointestinal tract harbors a complex microbial ecosystem that actively supports human health^1–3^ through beneficial functions that have co-evolved between the host and its resident microbiota^4–7^. Microbial persistence, defined here as the capacity of a population to remain detectable and maintain abundance during environmental perturbation^8^, is crucial to sustain these beneficial interactions. However, following initial colonization^8,9^, it is continuously challenged by host responses^10^, inter-bacterial competition^11^, and environmental fluctuations in the gut physical environment^4,12^. Maintaining stable populations in the face of these constraints is a central feature of microbial ecology, and reflects the combined effects of multiple species-specific physiological determinants, including growth, death, survival, and adaptive responses^13–17^.

Importantly, disentangling the individual contributions of these physiological determinants *in vivo* is challenging, in part due to the inherent complexity of native communities and methodological limitations^18–20^. To overcome these challenges, we seek to characterize intrinsic microbial phenotypes in response to physical perturbations by combining controlled *in vitro* assays, which isolate species-specific physiological responses, with *in vivo* models. Among physical perturbations relevant to the gut, osmotic stress may provide an experimentally tractable system in which intrinsic physiological responses can be isolated from more complex ecological interactions, particularly under severe conditions, a possibility we test directly below. In the gut, osmotic upshifts can be driven by a variety of factors, such as host malabsorption, disease, and the use of osmotic laxatives such as polyethylene glycol (PEG)^12,21^. Osmotic perturbations drastically reshape the gut ecosystem and microbiome, leading to taxon-specific responses which we hypothesize provide insights into *in vivo* persistence outcomes^22–24^. In previous work, we found that strain-specific behaviors and genomic stress-response genes were consistent with *in vivo* community resilience^25^, but it remains unknown whether intrinsic phenotypes can quantitatively predict *in vivo* community responses.

Here, we investigate which aspects of microbial physiology contribute to persistence *in vivo* under osmotic stress. We focus on growth and mortality as intrinsic determinants using *in vitro* assays and relate these measurements to *in vivo* abundance in a defined microbial community. We then examine whether adaptive responses during prolonged osmotic stress further contribute to persistence *in vitro*. We focus on *Bacteroides thetaiotaomicron*, a prominent gut symbiont, to investigate whether adaptation to osmotic stress through reduced mortality and genomic phase variation further contributes to persistence. Together, our findings identify growth capacity as a predictor of *in vivo* abundance during osmotic perturbation and show in *B. thetaiotaomicron* that reduced mortality provides an additional route to persistence under prolonged osmotic stress.

## Results

### Persistence *in vivo* is strongly associated with intrinsic growth capacity under osmotic stress

To investigate microbial persistence across a range of intestinal osmolalities, we modulated malabsorption in a gnotobiotic mouse model using polyethylene glycol (PEG), a well-characterized osmotic laxative^12,26^. Five PEG concentrations in drinking water generated a gradient of gut osmolality and motility (Fig. 1A, S1). Germ-free mice were colonized with a 10-member community^27^, comprising representatives of the five major bacterial phyla of the human gut microbiome (Bacteroidota, Bacillota, Actinomycetota, Pseudomonadota, Verrucomicrobiota) and spanning a wide range of osmotic tolerances^25^. Following community equilibration, the mice were exposed to PEG for 6 days before being euthanized for analysis. PEG treatment led to a range of cecal osmolalities from ∼350 to ∼850 mOsm/kg in 0 and 15% PEG, respectively (Fig. 1B). Increasing PEG concentrations progressively reduced community persistence, as evidenced by a dosage- and time-dependent decline in total bacterial abundance. Daily spot plating of fecal samples during PEG treatment revealed an approximate 10-fold reduction in total bacterial load in mice treated with 15% PEG (Fig. 1C), which was consistent with a decrease in cecal bacterial genomic load measured by species-specific droplet digital PCR (ddPCR) (Fig. 1D, S2, and Table S1). To measure the persistence of individual species, we performed shotgun sequencing on cecal DNA. We measured the Shannon diversity index across all mice, observing a PEG- and osmolality-dependent decrease in alpha diversity (Fig. 1E), driven by the depletion and disappearance of osmosensitive taxa, specifically *Muribaculum intestinale*, *Akkermansia muciniphila*, *Faecalibacterium prausnitzii, Agathobacter rectalis,* and *Bacteroides ovatus* (Fig. 1E). Conversely, increasing osmolality selected for osmotolerant taxa, with *Bacteroides thetaiotaomicron* and *Enterococcus faecalis* expanding to account for most of the community’s relative abundance under stress (Fig. 1E).

**FIGURE 1:**
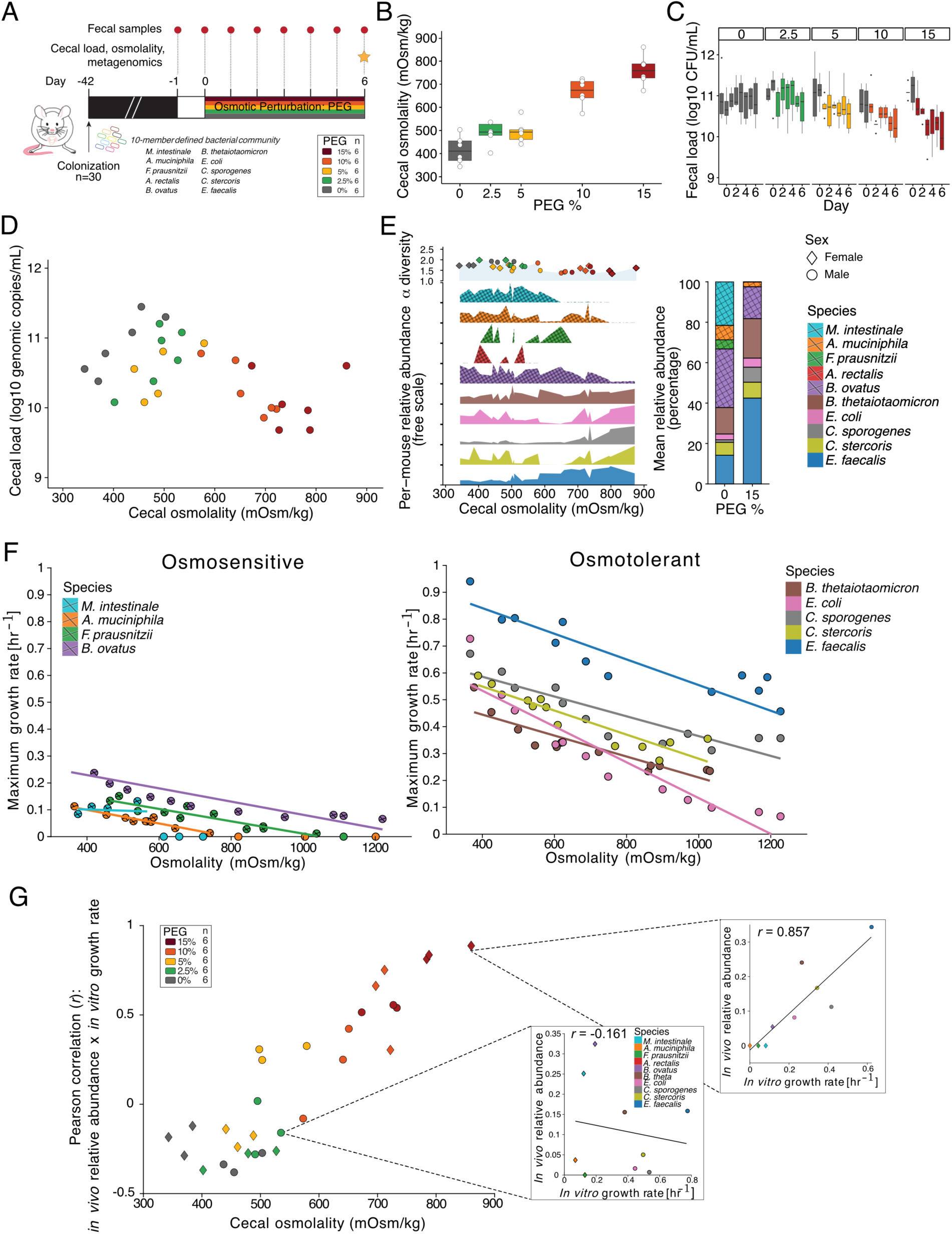
Inferred maximum growth rate *in vitro* is increasingly predictive of abundance *in vivo* at higher osmolality. **(A)** Experimental schematic and sample collection protocol. Germ-free mice (n=30) were colonised with 10 bacterial strains across five major bacterial phyla of the human gut microbiome (Bacteroidota, Bacillota, Actinomycetota, Pseudomonadota, Verrucomicrobiota) and allowed to equilibrate for 42 days, after which they were treated with 0%, 2.5%, 5%, 10%, 15% PEG in drinking water (shown by color) *ad libitum* for 6 days. **(B)** Cecal osmolality after 6 days of PEG treatment. Black-lined dots represent individual mice. **(C)** Total bacterial load of fecal pellets collected daily, measured by spot plating. Boxplots are color coded based on PEG dosage according to (A). Grey bars represent pre-treatment collection days or 0% PEG. **(D)** Total cecal bacterial load measured with species-specific droplet digital PCR (ddPCR) primers (Table S1) as the sum of genomic copy abundance of individual strains within each mouse (Methods and Fig. S2). Color represents PEG dosage. **(E)** Cecal relative abundance of individual species calculated from shotgun sequencing data across individual mice (bottom-left), averaged for 0% and 15% PEG mice (bottom-right) and alpha diversity (Shannon Index, top). In the lower panel, color represents bacteria while hatches indicate microbes whose relative abundance decreased in 15% PEG mice compared to 0% PEG. In the top panel, color represents PEG dosage and shape represents the sex of mice. **(F)** Species-specific linear regressions of the *in vitro* maximum growth rates across osmolalities in PEG-supplemented media for osmosensitive (left) and osmotolerant (right) community members. **(G)** Pearson correlation coefficient calculated from linear regressions between *in vivo* relative abundance and *in vitro* maximum growth rate estimated from (F) based on the cecal osmolality of each mouse. Each point represents a mouse where color is PEG dosage. The boxes contain 2 examples of how individual Pearson correlation coefficients were calculated across all 30 mice (see Fig. S3 for all mice).

**FIGURE 2:**
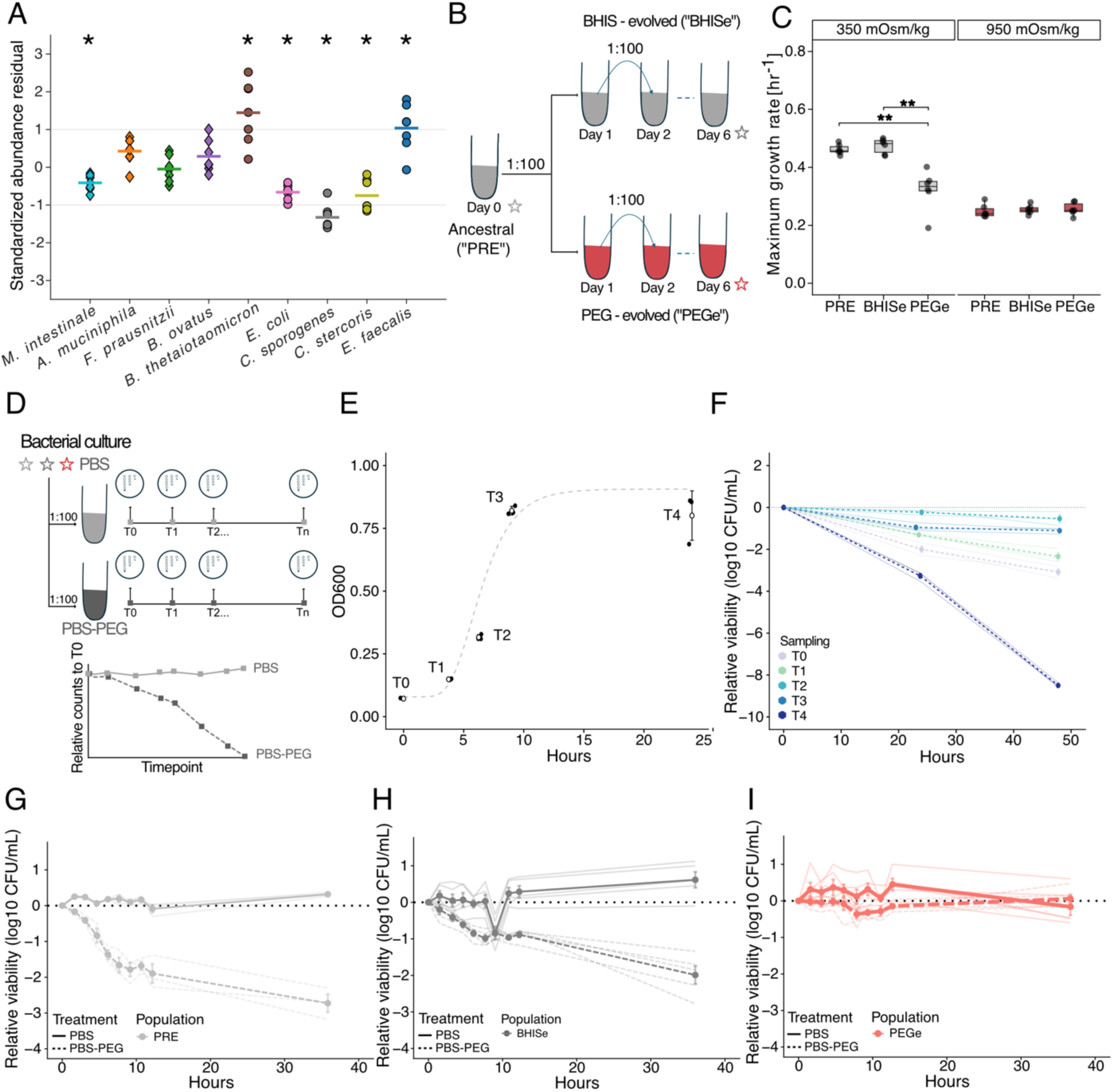
*Bacteroides thetaiotaomicron in vitro* mortality burden is reduced following prolonged exposure to PEG. **(A)** Species-specific residuals from the within-mouse linear regressions shown in Fig. 1G and Fig. S3, calculated as observed minus predicted *in vivo* relative abundance for mice with cecal osmolality >700 mOsm/kg (n=7). Predicted abundance was obtained from the regression of community relative abundance against species-specific predicted in vitro maximum growth rate within each mouse. Positive values therefore indicate species present at higher abundance than expected from the community-wide growth-abundance relationship. Statistical comparisons were performed using one-sample t-tests, followed by Bonferroni correction for multiple comparisons (*p < 0.05). **(B)** Experimental schematic of *in vitro* evolution assay. Biological triplicates of ancestral *B. thetaiotaomicron* (“PRE”) were sub-cultured daily for 6 days in either BHIS (“BHISe” in light grey, ∼350 mOsm/kg) or PEG-supplemented BHIS (“PEGe” in light red, ∼950 mOsm/kg). **(C)** Inferred maximum growth rate *i*n *vitro*, obtained by performing a least-squares fit to the Gompertz equation^28^, of *B. thetaiotaomicron* PRE, BHISe, and PEGe exponential cells cultured in BHIS (∼350 mOsm/kg, left) and BHIS-PEG media (∼950 mOsm/kg, right) (n=6). Statistical significance was determined by a two-tailed, paired t-test (**p < 0.01). **(D)** Experimental schematic of the mortality burden assay. Following passaging (B), exponential stage PRE, BHISe, and PEGe populations were diluted 1:100 into either baseline PBS (light grey, ∼300 mOsm/kg) or PEG-supplemented PBS (dark grey, ∼950 mOsm/kg). Samples were collected regularly and spot plated to obtain longitudinal live counts relative to the first timepoint T0. **(E)** *B. thetaiotaomicron* PRE samples (n=3) were collected at 5 distinct timepoints, labelled T0 to T4 with a Gompertz curve fitted in the dashed line. **(F)** At each timepoint, *B. thetaiotaomicron* was exposed to PBS-PEG (∼950 mOsm/kg, dashed lines) and spot plated to investigate the impact of growth stage on mortality burden. Color represents the sampling time, thin lines the individual biological replicates, and thick lines the mean mortality. **(G-I)** *In vitro* mortality burden of PRE (G), BHISe (H), and PEGe (I) *B. thetaiotaomicron* populations (n=3) in either PBS (∼350 mOsm/kg, solid lines) or PBS-PEG (∼950 mOsm/kg, dashed lines). The dotted line at y = 0 is used as a visual reference for T0.

To determine whether intrinsic physiological performance could explain differences in persistence across this osmotic gradient, we quantified species-specific growth responses across a finely resolved range of PEG-induced osmolalities using monocultures *in vitro* (Methods). This profiling revealed a clear division in growth capacity consistent with the survival patterns observed *in vivo* (Fig. 1F). Specifically, osmosensitive species that were severely depleted at high PEG *in vivo* showed lower maximum growth rate across all conditions tested, including at baseline, compared to osmotolerant species (Fig. 1F). To investigate whether *in vitro* population growth dynamics were sufficient to predict *in vivo* persistence, we correlated the *in vivo* abundance of each species at a specific osmolality with its *in vitro* maximum growth rate estimated at the corresponding osmolality (Fig. 1G). At low osmolalities, *in vitro* maximum growth rates were modestly anti-correlated with *in vivo* abundance (r ∼ -0.3) (Fig. 1G), suggesting that conditions of low perturbation may select for relatively slower growing species but that factors other than osmotic tolerance shape species abundance and persistence. However, the relationship between *in vitro* growth and *in vivo* persistence strengthened progressively with increasing osmolality, reaching r ∼ 0.9 at ∼850 mOsm/kg (Fig. 1G and S3). These findings indicate that as intestinal osmolality increases, community composition increasingly favors species that can sustain high growth rates under osmotic stress. These results were consistent when absolute abundances measured via ddPCR were considered rather than compositional data (Fig. S4 and S5). Altogether, these data show that osmotic stress drives major microbial community restructuring and that species-specific *in vitro* maximum growth rates may be leveraged to predict microbial persistence *in vivo* under increasingly perturbed environments.

### Mitigation of mortality burden is a key adaptive strategy for *B. thetaiotaomicron* persistence under osmotic stress

Given the close correlation between abundance in vivo and growth rates in vitro at high osmolality, we next asked whether individual species showed systematic deviations from the community-wide relationship used in Fig. 1G. We therefore quantified, for each species, the difference between its observed *in vivo* relative abundance and the abundance expected from the within-mouse linear relationship between growth rate and community abundance for mice with cecal osmolalities >700 mOsm/kg (Fig. 2A). The more osmosensitive species *A. muciniphila*, *F. prausnitzii*, and *B. ovatus* showed little systematic deviation from this relationship, whereas the more osmotolerant members *B. thetaiotaomicron* and *E. faecalis* were consistently observed at higher abundances than expected from population growth rate alone. We therefore used these residuals as a diagnostic to identify species whose abundance was not fully captured by the growth-abundance relationship, rather than as an independent predictive test.

To investigate the potential contribution of survival and adaptation, we focused on *B. thetaiotaomicron* as a model for studying physiological mechanisms that contribute to persistence under osmotic stress. To simulate the prolonged PEG osmotic stress exposure of the *in vivo* model, we passaged ancestral cells (“PRE”) for 6 days in either standard BHIS (“BHISe”, 350 mOsm/Kg) or PEG-supplemented BHIS media (“PEGe”, 950 mOsm/Kg) (Fig. 2B). We detected significantly lower inferred maximum growth rate in baseline BHIS media among evolved PEGe cells compared to PRE and BHISe controls, but no statistical differences under high osmolality medium (Fig. 2C), indicating that while PEGe cells may have undergone physiological changes, adaptation was not associated with enhanced growth capacity.

We then reasoned that, while inferred maximum growth rates were unchanged compared to controls following adaptation, population growth reflects the net balance of cellular replication and cell loss and therefore does not reveal their individual contributions to persistence under high osmolality. We thus sought to directly quantify survival via relative viability, reasoning that persistence under high osmolality may depend not only on growth capacity but also on reduced cell loss, particularly under conditions of limited growth. We therefore characterised mortality burden using a nutrient-restricted cell viability assay (Methods), measuring the decline in viable cell counts during exposure to normal and high-osmolality PEG conditions in saline solution (PBS, Fig. 2D). These nutrient-restricted conditions were used to isolate differences in cell loss in the absence of population growth, rather than to directly recapitulate the intestinal environment. We first investigated how growth phase impacts mortality using ancestral cells exposed to PEG-supplemented PBS (PBS-PEG, Fig. 2E). We observed that survival depended strongly on physiological stage, with cells from early-, mid-, and late-exponential growth phases exhibiting substantially lower mortality burdens than late-stationary phase cells after 48 hours (log_10_ -2.3, -0.5, -1.1, respectively), whereas late-stationary phase cells exhibited substantially greater mortality (log_10_ -8.5) (Fig. 2F). To minimize growth-stage variability, all subsequent mortality assays were performed using mid-exponential cultures, which provide a reproducible physiological state across cultures. We then compared mortality burdens among the PRE, BHISe, and PEGe lineages. Under high-osmolality conditions, PRE and BHISe lineages exhibited a log_10_ reduction of -2.7 and -2.0, respectively, after 36 hours, compared to no death in the PBS control conditions (Fig. 2G, H). In contrast, the adapted PEGe lineages showed no differences in normalised log10 mortality in osmotic conditions compared to PBS control (0.1 and -0.2, respectively) (Fig. 2I), suggesting that prolonged exposure to high osmolality selected for a phenotype with markedly reduced mortality burden (Fig. 2I).

### Phase-variable loci are associated with *B. thetaiotaomicron* adaptation to osmotic stress

Having observed a reduction in mortality burden in PEG-adapted *B. thetaiotaomicron*, we next aimed to determine the genetic basis of this adaptive osmotolerant phenotype. To achieve this, we performed shotgun sequencing on PRE, BHISe, and PEGe populations (Methods). While we did not find fixed mutations, indels, or substitutions unique to PEG-evolved populations (Table S2), we identified regions with significant changes in phase orientation (Methods). Phase variation enables *Bacteroides* to reversibly alter gene expression through DNA inversions, commonly by switching the direction of promoters, generating phenotypic diversity that facilitates rapid adaptation to changing environmental conditions^29,30^. Specifically, we identified two intergenic invertons that exhibited significantly greater frequencies of the reverse orientation following PEG adaptation (Fig. S6). These invertible regions mapped between 1) BT0375 and BT0376, near the capsular polysaccharide 1 cluster (CPS1, between BT0376 and BT0400), and 2) BT4299 and BT4300, near the polysaccharide utilization locus 78/80 operon (PUL78/80, between BT4294 and BT4299). CPS clusters regulate the production of surface capsular polysaccharides, and they have been implicated in host immune evasion or modulation, host colonization, competitive fitness, and bacteriophage susceptibility^31,32^. Meanwhile, polysaccharide utilization loci such as PUL78/80 have been shown to encode outer-membrane complexes dedicated to the binding, internalization, and degradation of environmental carbohydrates, and are implicated in population persistence as well as interactions with other microbes^33,34^. Quantification of the CPS1 and PUL78/80 phase orientations following PEG adaptation showed consistent enrichment across independently passaged lineages, with the BT0376-ON conformation increasing to 35% in PEGe from 10% in PRE and 17% in BHISe (p < 0.001 and 0.01, respectively) (Fig. 3A). Similarly, the BT4299-OFF conformation rose significantly to 65% in PEGe, compared to 40% in PRE and 45% in BHISe (p < 0.01) (Fig. 3A). Together, these results suggest that osmotic adaptation is associated with coordinated changes in the orientation of specific phase-variable loci, consistent with reversible regulatory adaptation in the absence of fixed genetic mutations.

**FIGURE 3:**
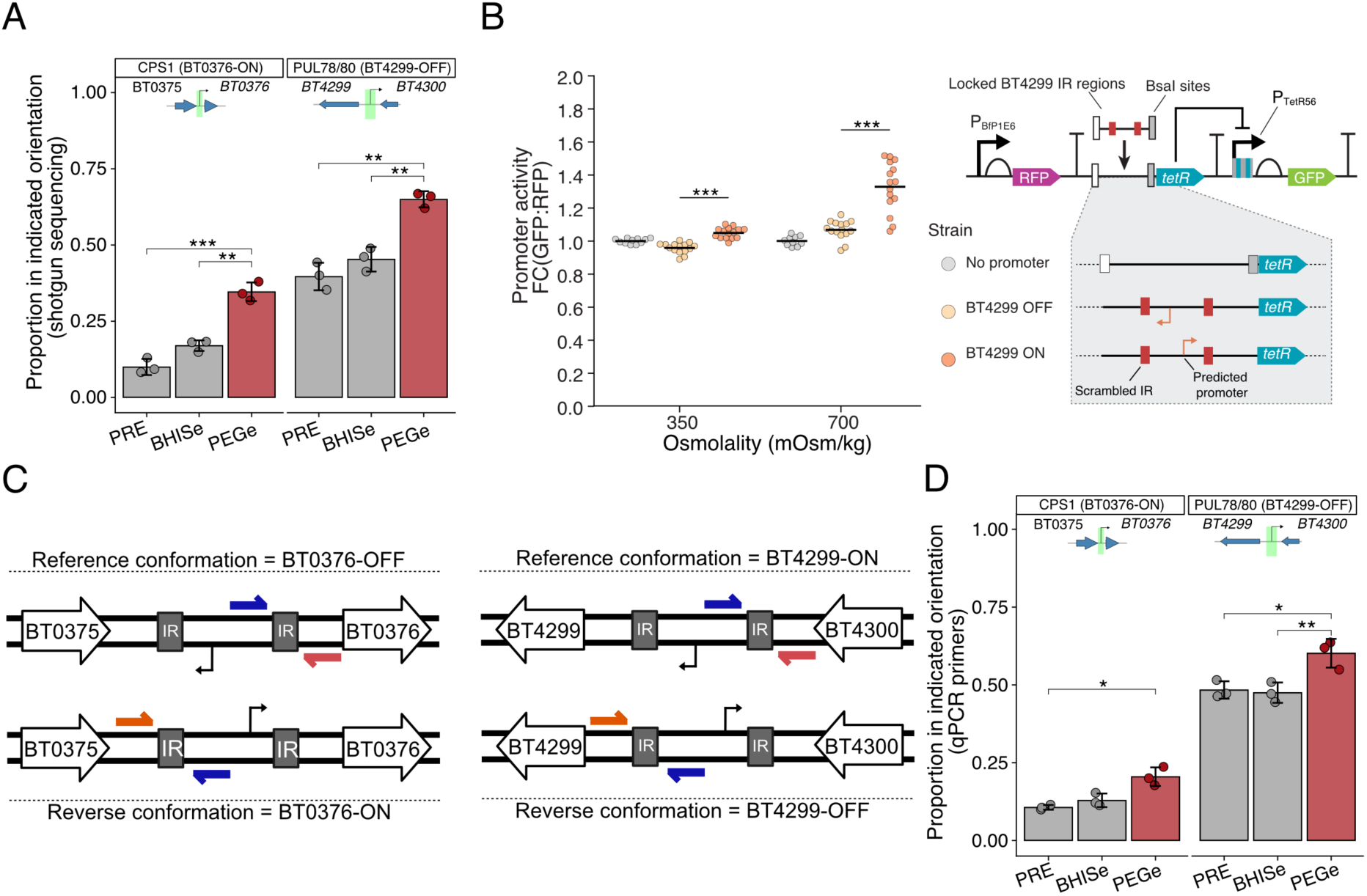
Prolonged exposure to PEG leads to altered phase variation in CPS1 and PUL78/80 invertons. **(A)** Proportion of *B. thetaiotaomicron* cells with invertible promoters in the reverse conformation for CPS1 (left) and PUL78/80 (right), respectively indicated as BT0376-ON and BT4299-OFF from shotgun sequencing on the PRE, BHISe, and PEGe populations (star in Figure 2B). Color indicates the osmolality of the passaging medium. The gene map schematic at the top of each facet indicates the location and directions of the invertons. Data points represent individual biological replicates (n=3), with large circles indicating the mean and error bars the standard deviation. Statistical significance was determined by a two-tailed, paired t-test (*p < 0.05, **p < 0.01, ***p < 0.001). **(B)** The upstream region of BT4299, containing the sequence between the two inverted repeats (IRs) locked in either of its two possible orientations was cloned into a reporter plasmid^35^ to measure promoter activity as the fold change (FC) in the GFP to RFP ratio (FC[GFP:RFP]) relative to a no promoter control (see Table S3 and S4). Activation of the candidate constructs was measured during growth at baseline (350 mOsm/kg) or 700 mOsm/kg media. Statistics were calculated using an unpaired t-test comparing the two constructs. (***p < 0.001). **(C)** Schematic of the reference (top) and reverse (bottom) promoter conformations in both CPS1 (left) and PUL78/80 (right) intergenic invertons. qPCR primers were designed and validated to detect cells with promoters in both orientations, using fixed primers outside the IRs, colored orange and red, and a primer within the invertible region, colored blue (Table S1). **(D)** Proportion of *B. thetaiotaomicron* cells with invertible promoters in the reverse conformation for CPS1 (left) and PUL78/80 (right). Ratios were determined using the qPCR primers outlined in (C) on the same DNA samples sent for shotgun sequencing and used in (A). Statistics were calculated and displayed as stated in (A).

While the CPS1 invertible promoter location and direction have been characterized^31^, we sought to validate the orientation of the regulatory element within the PUL78/80 inverton in *B. thetaiotaomicron*. To achieve this, we engineered a dual-fluorescent reporter system with the invertible region locked in either the ON or OFF conformation (Fig. 3B, Table S3 and S4)^35^. We found that the PUL78/80 inverton contains an invertible promoter that is active in the reference (BT4299-ON) orientation. Under baseline osmolality, the “ON” construct exhibited significantly higher promoter activity than the “OFF” construct (p < 0.001), which conversely generated background levels of activation similar to a no-promoter control (FC ∼ 1.0). The promoter was also active in the “ON” conformation at 700 mOsm/kg, compared to low or no activity in the “OFF” and no-promoter constructs (p < 0.001) (Fig. 3B). These data confirmed that the PUL78/80 inverton functions as an invertible promoter, enabling interpretation of phase orientation in subsequent experiments.

To streamline the detection of these structural variants across numerous samples without sequencing, we designed orientation-specific qPCR primers (Fig. 3C, Table S1). This qPCR assay reliably recapitulated the phase-orientation changes, capturing the same enrichment patterns across biological replicates, including enrichment of the BT0376-ON conformation in PEGe cells compared to PRE, as well as significant shifts toward the BT4299-OFF conformation relative to PRE and BHISe (p < 0.05) (Fig. 3D). We therefore used orientation-specific qPCR as a robust, sequencing-independent method to quantify phase switching at both loci in subsequent experiments.

Altogether, these results identify two phase-variable loci associated with the adaptive response of *B. thetaiotaomicron* to osmotic stress and establish orientation-specific qPCR as a robust method for monitoring their switching dynamics.

### Selection for the OFF orientation of the phase-variable PUL78/80 locus correlates with *B. thetaiotaomicron* survival under osmotic stress

Given the established role of phase variation in stress responses, we sought to further characterize the stability and reversibility of the CPS1 and PUL78/80 phase-state distributions under changing osmotic conditions. To do so, we conducted a second 6-day passaging round of PRE, BHISe, and PEGe grown back into either BHIS or BHIS-PEG, where we expected a return to pre-stress ratios in PEGe cells cultured in BHIS and possibly a further increase in reverse conformation following the 6 additional days in BHIS-PEG (Fig. 4A). We found that the population distributions of both CPS1 and PUL78/80 phase variants reproducibly converged across independent lineages to stable, environment-specific equilibria regardless of evolutionary history (Fig. 4B). For example, when adapted PEGe lineages were returned to baseline media (PEGe-BHISe), the proportion of the BT4299-OFF conformation decreased from ∼60% down to ∼50%, effectively reverting to ancestral baseline levels (∼48%). Furthermore, consecutive passaging in PEG (PEGe-PEGe) did not significantly increase the reverse conformation ratio beyond a single round of selection (PEGe) (64% vs. 60%, respectively), suggesting that phase switching reaches optimal levels within 6 days of passaging. Together, these results indicate that osmotic stress establishes reproducible, environment-dependent phase-state equilibria rather than driving irreversible selection of specific variants.

**FIGURE 4:**
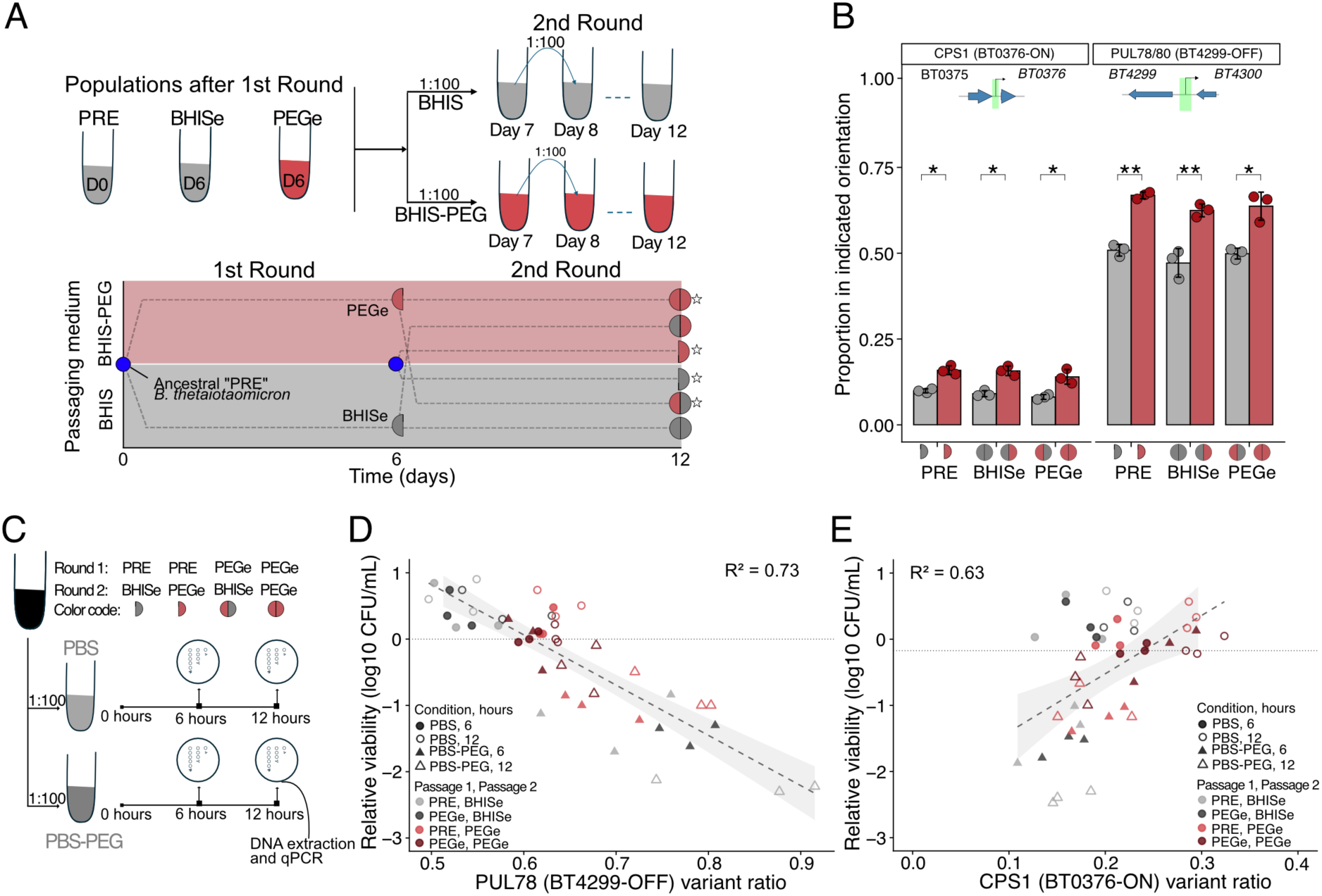
Phase variant switching is not affected by evolutionary history and specific orientations are enriched in surviving cells under PEG-induced osmotic stress. **(A)** Experimental schematic of the second round of passaging *in vitro* (top) and the overview of populations created (bottom). *B. thetaiotaomicron* PRE, BHISe, and PEGe populations from the first passaging round (1^st^ round, Figure 2B) were passaged following the same protocol (2^nd^ round). The ancestral *B. thetaiotaomicron* (PRE, in blue) is used to start all populations. Half circles represent populations that were passaged once. Grey indicates passaging in BHIS medium (∼350 mOsm/kg) and red passaging in BHIS-PEG (∼950 mOsm/kg). Full circles represent populations that have undergone two 6-day passaging events, where the left half of the circle represents the medium of the first round, and the right half the medium from the second round. **(B)** Proportion of *B. thetaiotaomicron* cells with invertible promoters in the reverse conformation for CPS1 (left) and PUL78/80 (right) (n=3). Ratios were determined using the qPCR primers outlined in Figure 3C on the populations obtained following the 2^nd^ round of passaging. The passaging history of each population is visually summarised using solid or split circles on the x-axis. The gene map schematic at the top of each facet indicates the location and direction of the invertons. Data points represent individual biological replicates (n=3), with large circles indicating the mean and error bars the standard deviation. Statistical significance was determined by a two-tailed, paired t-test (*p < 0.05, **p < 0.01). **(C)** Experimental schematic of the mortality burden assay performed to characterise the proportion of PUL78/80 **(D)** and CPS1 **(E)** invertons in surviving *B. thetaiotaomicron* cells across selected populations, also labelled with stars in (A). The dashed line and gray shaded arrow indicate a linear regression fitted on all data points.

Next, we hypothesized that specific phase variants may confer a survival advantage during osmotic stress and would therefore become enriched following acute osmotic challenge. To investigate this, we subjected select populations to the nutrient-restricted mortality assay and quantified phase variant reverse conformation ratios among surviving cells (Fig. 4C). We found that osmotic shock strongly enriched the BT4299-OFF conformation in the PUL78/80 phase variant (Fig. 4D), consistent with its enrichment after prolonged exposure to PEG (Fig. 3D). The frequency of the BT4299-OFF orientation increased progressively with increasing mortality burden (R² = 0.73). Specifically, the BT4299-OFF orientation increased to ∼90% among surviving cells in PRE-BHISe populations that experienced approximately ∼100-fold depletion during osmotic shock (Fig. 4D). Conversely, adapted PEGe-PEGe populations that already exhibited low mortality during osmotic shock maintained BT4299-OFF ratios nearly identical to their pre-assay baselines (∼60%). Unlike PUL78/80, the CPS1 phase variant showed a weaker pattern, with the BT0376-ON proportion slightly decreasing as viability decreased, thus indicating a small enrichment in BT0376-OFF cells (Fig. 4E). These results suggest that CPS1 phase orientation is less strongly associated with survival under osmotic stress. In particular, the selective enrichment of the BT4299-OFF conformation following acute osmotic shock identifies the PUL78/80 inverton as a strong candidate mediator of enhanced survival.

### Rapid phase switching is coupled to transcriptional remodeling of CPS1 and PUL78/80

Although PUL78/80 showed the stronger association with survival, both PUL78/80 and CPS1 underwent reproducible phase switching during prolonged osmotic stress. We therefore asked how rapidly these phase-state changes occurred. We alternated growth of PRE and PEGe populations between BHIS or BHIS-PEG media and sampled repeatedly throughout the time course. We found that *B. thetaiotaomicron* exhibited asymmetric inversion dynamics following the addition or removal of osmotic shock. Specifically, the transition to the reverse orientation under high osmolality in PRE cells was relatively rapid, with the fraction of BT0376-ON cells increasing from ∼15% to ∼23% and BT4299-OFF cells from 52% to 80% within 30 hours (Fig. 5A, B). Conversely, reverse orientations persisted substantially longer after populations were returned to BHIS. Despite a gradual decline in both BT0376-ON and BT4299-OFF ratios over 3 days in BHIS, neither population returned to ancestral baseline levels within the experimental window (Fig. 5A, B). Together, these temporal dynamics demonstrate that *B. thetaiotaomicron* rapidly adopts phase-state distributions characteristic of high osmolality conditions, while retaining them long after the stress is removed, suggesting that phase variation provides a sustained physiological memory of previous osmotic exposure.

**FIGURE 5:**
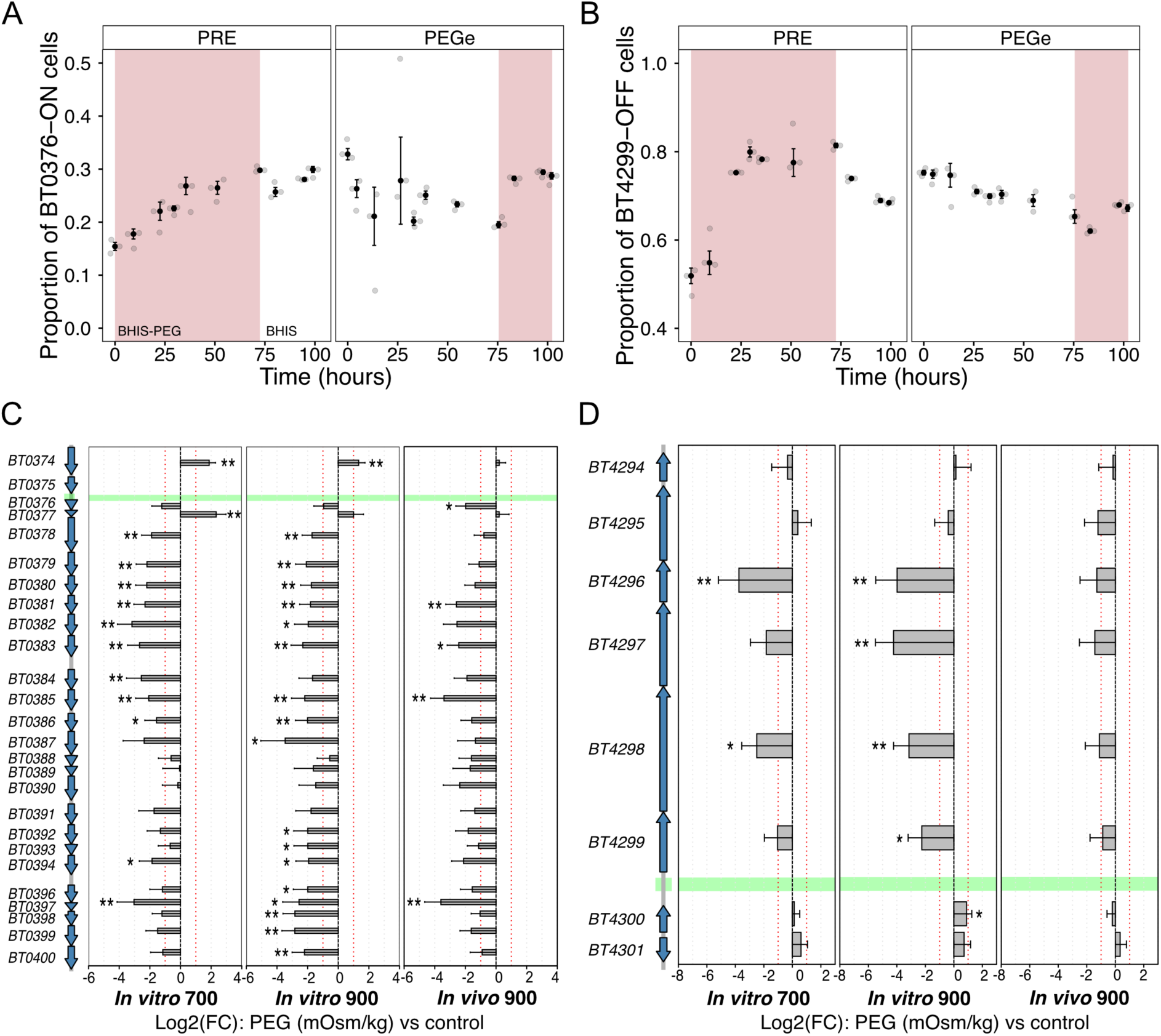
CPS1 and PUL78/80 phase switching following PEG exposure begins within a day and is associated with transcriptional changes. (A-B) Temporal dynamics of phase switching in the CPS1 (A) and PUL78/80 (B) intergenic invertons starting from ancestral *B. thetaiotaomicron* PRE (left) and PEGe (right) populations. Red shading indicates growth in BHIS-PEG medium (∼950 mOsm/kg) while no shading indicates growth on baseline BHIS medium (∼350 mOsm/kg). Jittered grey points indicate independent biological replicates (n=3) and black points represent the mean of each group. Error bars indicate the standard deviation. **(C-D)** Transcriptional changes of the CPS1 (C) and PUL78/80 (right) gene clusters from published data in McCallum et al. (2026)^35^. On the left are the genomic maps showing the CPS1 loci BT0376-0400 (25 loci) and the PUL78/80 loci BT4294-4299 (6 loci) together with adjacent genes, respectively BT0374-BT0375 and BT4300-BT4301. On the right are the log2 fold changes relative to baseline controls for *in vitro* PEG (700 and 900 mOsm/kg) and *in vivo* PEG (900 mOsm/kg) conditions. The green horizontal bar indicates the approximate location of the invertible promoters. Asterisks denote significant differential expression (*p < 0.05, **p < 0.01).

Given the rapid phase switching under osmotic shock, we next asked whether these changes were associated with altered transcription of the CPS1 and PUL78/80 loci. We leveraged an existing transcriptomic dataset profiling *B. thetaiotaomicron* after 6-8 hours of exposure to three PEG hyperosmotic conditions: *in vitro* (700 and 900 mOsm/Kg) and *in vivo* (900 mOsm/Kg) compared to baseline^35^ (Fig. 5C, D). We observed broad transcriptional repression across genes downstream of both phase variants, consistent with the increased frequencies of the reverse conformations observed in our assays. Specifically, we observed significant transcriptional repression of the majority of the CPS1 genes (BT0376 to BT0400, i.e., 25 loci) across all conditions, except for the UpxZ anti-terminator antagonist (BT0377) which was upregulated (Fig. 5C). These results are consistent with the predicted transcriptional consequences of the BT0376-ON phase-state distribution, where the UpxZ (BT0377) represses the transcription anti-terminator UpxY (BT0376), thus down-regulating the entire cluster.

Similarly, consistent with enrichment of the BT4299-OFF conformation, which directs the promoter away from the downstream operon, the dataset showed strong repression of the downstream PUL78/80 genes (the six loci BT4294-BT4299), alongside modest upregulation of BT4300 and BT4301 (Fig. 5D). Together, these transcriptomic data show that osmotic-shock-induced phase switching is accompanied by coordinated transcriptional remodeling of the CPS1 and PUL78/80 loci, consistent with the predicted regulatory consequences of these inversion events.

### Stress-associated PUL78/80, but not CPS1 phase states are maintained *in vivo*

To determine whether phase-variable states identified during osmotic adaptation also occurred during gut colonization, we quantified the orientation of the CPS1 and PUL78/80 invertons using selected cecal DNA samples from the mouse experiment in Figure 1. We quantified phase orientation using the qPCR assay described in Fig. 3C, as insufficient read coverage in our short-read sequencing dataset precluded reliable estimation of inversion orientation. The CPS1 orientation varied with intestinal osmolality, shifting from values comparable to evolved populations at lower osmolality toward the pre-evolved state at higher osmolality (Fig. 6A). In contrast, the PUL78/80 locus was strongly enriched in the OFF orientation across all conditions tested, exceeding the fraction observed in both pre-evolved and evolved populations (Fig. 6B). Notably, the ratio of BT0376-ON cells decreased to nearly 0% in the 5% PEG dosage at ∼600 mOsm/kg before increasing in the higher PEG dosage samples. These results indicate that the phase-variable PUL78/80 states identified during experimental evolution are also represented *in vivo*, while CPS1 states vary strongly based on the environment.

**FIGURE 6:**
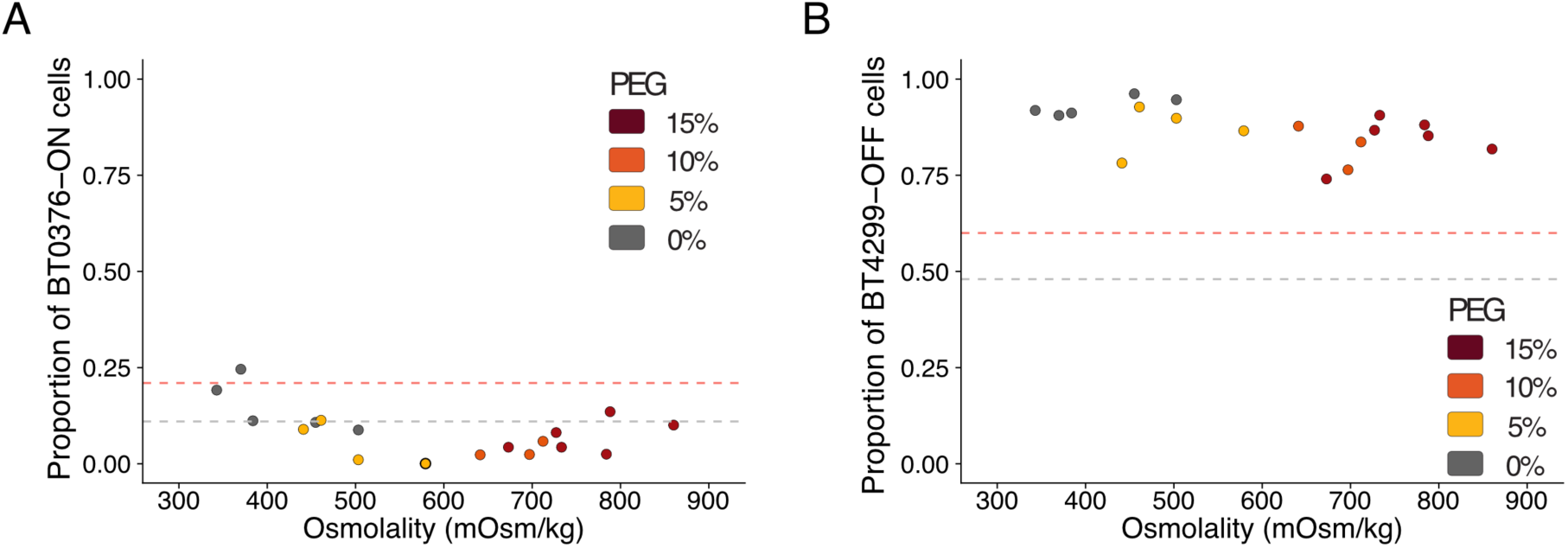
Stress-associated PUL78/80 but not CPS1 phase orientations are observed *in vivo*. **(A)** Proportion of BT0376 (CPS1) ON cells measured with qPCR primers in mice from the experiment shown in Fig. 1 as a function of intestinal osmolality. **(B)** Proportion of BT4299 (PUL78/80) OFF cells measured in the same mice as a function of intestinal osmolality. Points represent individual mice and were selected among all *in vivo* samples to account for the full range of cecal osmolalities. Color represents the PEG concentration used to modulate intestinal osmolality. The red and gray dashed lines indicate the mean proportion of each phase state measured in the PEG-evolved (PEGe) and pre-evolved (PRE) populations, respectively, following prolonged *in vitro* osmotic stress.

## Discussion

Predicting microbial persistence *in vivo* requires deconvoluting their complex interactions with other microbes, the host, and host environment. However, severe environmental perturbations can establish limiting conditions that become major drivers of community composition^12,25^, enabling investigations of intrinsic microbial phenotypes as predictors of *in vivo* abundance. In this study, we used a defined microbial consortium to demonstrate that intrinsic *in vitro* maximum growth rate strongly correlates with *in vivo* relative and absolute abundance under severe osmotic stress, corroborating the importance of growth as a primary determinant of persistence *in vivo*. Importantly, the higher-than-predicted *in vivo* abundance of *B. thetaiotaomicron* and *E. faecalis* indicated that *in vitro* growth capacity alone could not fully account for persistence, suggesting that additional physiological factors influence survival under osmotic stress. We therefore investigated adaptive responses to prolonged osmotic stress in *B. thetaiotaomicron* and found that adaptation was associated with a marked reduction in mortality burden, identifying survival during osmotic stress as an important contributor to persistence under these conditions. This process was associated with genomic phase variation at the PUL78/80 inverton. Our data show that phase variation in *B. thetaiotaomicron* under osmotic stress occurs rapidly and reversibly, independently of evolutionary history, and is correlated with repression of both clusters, congruent with the switch in phase orientation. Together, these findings highlight the potential of using intrinsic microbial phenotypes as predictors of *in vivo* abundance, while also identifying reduced mortality and associated phase variation, particularly at PUL78/80, as key features of adaptation in *B. thetaiotaomicron* during osmotic perturbations.

Environmental disturbances, including osmotic perturbations, lead to major changes in microbial community composition^12,36^ and increased susceptibility to pathogens^24,37^. Under these limiting conditions, intrinsic microbial growth traits have been repeatedly linked to restructuring outcomes, with fast growth being a key determinant of persistence both *in vivo* and during high mortality in pairwise competition^13,37,38^. Osmotic perturbations further highlight the importance of fast growth by increasing luminal liquidity and flow rate, which impose dynamic minimum growth thresholds to avoid mechanical washout^14,39^. Consistent with these observations, our results confirm the central role of growth capacity during osmotic perturbations and demonstrate that intrinsic maximum growth rate predicts *in vivo* abundance under osmotic stress, extending previous observations that *in vitro* stress tolerance predicts bacterial responses to environmental perturbations *in vivo*^25^ (Fig. 1G). Notably, predictions based on absolute and relative abundance were comparable in our defined consortium, although this correspondence may diminish in more complex communities due to the challenges associated with quantifying individual members (Fig. S4).

While intrinsic growth capacity strongly predicts persistence during osmotic perturbations, recurrent exposure to these conditions also creates selective pressure for adaptive responses that further enhance survival. Gut microbial communities experience frequent and potentially long-lasting environmental perturbations, creating conditions where resilient members may develop adaptations to overcome recurrent physical challenges. Consistent with prior research in *E. coli*^40^, *B. thetaiotaomicron* adapted to high osmolality displayed growth defects in baseline media, yet maintained equivalent maximum net growth rates under osmotic stress compared to unadapted populations (Fig. 2C). In addition, adapted *B. thetaiotaomicron* showed reduced mortality burden in high osmolality (Fig. 2I) compared to controls (Fig. 2G,H). Because net population growth reflects the balance between cellular replication and cell loss, the combination of unchanged maximum net growth and reduced mortality raises the possibility that adaptation is accompanied by reduced replication under stress. A shift toward lower replication and mortality rates could represent a fitness trade-off that promotes persistence during prolonged osmotic stress. This interpretation is consistent with studies in *E. coli*, where slower growth is accompanied by exponentially slower death under starvation^41^. This behavior may also apply to osmotic stress as cells undergo similar physiological changes under nutrient deprivation and osmotic challenges^42^ and may help explain why some gut commensals persist during physical perturbation without maximizing growth.

Given the frequent and transient nature of these gut perturbations, short-term adaptation may favor rapid and reversible genetic mechanisms. Consistent with this, we observed significant phase switching in the PUL78/80 and CPS1 clusters of *B. thetaiotaomicron* evolved in high osmolality (Fig. 3A,D). Phase variation is a well-documented reversible adaptive strategy during environmental perturbations, including osmotic stress and antibiotic exposure^30,43,44^. Upon exposure to high osmolality, both clusters show a rapid and significant shift towards the reverse conformations (Fig. 5A,B), corresponding to a greater proportion of cells with PUL78/80 OFF and CPS1 ON states. However, this transition did not reach fixation within the population, and the removal of the stressor resulted in a distinctly slower return to the baseline forward conformations (Fig. 5A,B). This asymmetric switching dynamic is consistent with a bet-hedging strategy that maintains a subpopulation primed for repeated exposure to osmotic stress. Notably, the physical shift to these reverse conformations directly correlated with the transcriptional regulation of both the PUL78/80 and CPS1 clusters (Fig. 5C,D) and may relate to the selection of PUL78/80 OFF under high osmotic stress (Fig. 4D). Because both loci encode components of the bacterial outer surface, these phase variation and transcriptional data suggest that remodeling cell-envelope-associated functions may contribute to osmotic stress adaptation in *B. thetaiotaomicron*.

Consistent with the physiological relevance of these adaptive phase states, we found that the stress-associated PUL78/80 OFF orientation was enriched across the full range of intestinal osmolalities examined *in vivo*, whereas CPS1 exhibited a more variable pattern (Fig. 6). This finding suggests that the PUL78/80 OFF orientation may represent a broader adaptation to the intestinal environment rather than a specific response to elevated osmolality, potentially reflecting selection imposed by other host-associated stresses that favor maintenance of this phase state. Similarly, selection acting on CPS1 *in vivo* may be shaped by additional host-derived pressures. Previous studies have shown that individual *B. thetaiotaomicron* capsules differentially regulate host immune responses, with CPS1 eliciting distinct IgA and T-cell responses compared with acapsular or alternative capsule variants^45,46^. These observations raise the possibility that immune-mediated selection constrains CPS1 phase variation *in vivo*, potentially offsetting selection for osmotic stress-associated phase states. Together, these findings suggest that osmotic stress is one of multiple environmental pressures shaping phase variation *in vivo*, providing new insight into the regulatory landscape governing these adaptive responses.

In conclusion, our findings demonstrate that intrinsic growth capacity can be leveraged to predict *in vivo* persistence during osmotic perturbations. Our data also identify mitigation of mortality burden as an important determinant of persistence in *B. thetaiotaomicron*, with PUL78/80 phase variation strongly associated with survival under osmotic stress. Future studies aimed at defining the mechanisms underlying persistence during osmotic stress will further refine the quantitative framework established in this study, while clarifying how phase-variable adaptation contributes to survival under osmotic stress. Ultimately, integrating physical growth constraints with these discrete adaptive mechanisms will improve our ability to predict microbiome stability and engineer targeted interventions during recurrent environmental disruptions.

### Limitations of the study

The results presented in this study include several limitations. First, further characterizing the full landscape of phase variation will refine the framework presented here. Current short-read sequencing restricts detection to intergenic phase variants; thus applying long-read methodologies will enable the detection of intragenic, partially intragenic, and epigenetic modifications^47,48^. Resolving these additional layers of phase variation will provide a more comprehensive understanding of how diverse phase-variable strategies regulate adaptation during physical perturbations. Beyond this, extending the quantitative framework presented in this work to more complex yet defined microbiomes, such as the hCom synthetic communities^19^, will provide an important test of the predictability of intrinsic growth capacity. Because *in vitro* maximum growth rate successfully predicted both absolute and relative abundance, applying high-resolution shotgun sequencing to these defined communities should enable robust evaluation of this framework without requiring absolute quantification.

## Resource availability

### Contact for Reagent and Resource Sharing

Further information and requests for resources and reagents may be directed to and will be fulfilled by the lead contact, Carolina Tropini.

## Materials availability

All plasmids generated in this study can be made available from the lead contact upon reasonable request.

## Acknowledgments

The authors acknowledge that the land we performed this research on is the traditional, ancestral, and unceded territory of the xwməθkwəy̓əm (Musqueam) Nation. We encourage others to learn more about the native lands in which they live and work at https://native-land.ca/

H.G. would like to thank Paula von Sperling, Ian Ghezzi, G.V., B.B., and L.V. for the support throughout the development of this work. We also thank Angele Arrieta for research support, Natalia Carranza Garcia for support with mouse work, and the W. Garfield Weston Foundation / Weston Family Microbiome Initiative. This work received support from Gut4Health (RRID:SCR_023673) at the BC Children’s hospital Research Institute, and the Sequencing + Bioinformatics Consortium (SBC) and Advanced Research Computing (ARC) at the University of British Columbia. The authors acknowledge support from 4-Year Fellowship (to H.G. and J.C.B.), Michael Smith Health Research BC Trainee Award (RT-2023-3174, to M.H.), Canadian Institute for Advanced Research/Humans and the Microbiome (FL-001253 Appt 3362, to C.T.; CF-0341 - CP24-007 to N. G.-Z. and C.T.), Michael Smith Foundation for Health Research Scholar Award (18239, to C.T.), Johnson and Johnson Women in STEM2D Award (015007, to C.T.), Canada Foundation for Innovation/Infrastructure Operating Fund (38277, to C.T.), Canadian Institutes of Health Research (CIHR) Priority Announcement: Infection and Immunity (PTT 190382, to C.T.), US National Institutes of Health National Institute of General Medical Sciences award (R35GM151023, to N.G.), National Science Foundation CAREER award (no. 2240098, to N.G.), Paul Allen Research Foundation grant (to N.G.).

## Author contributions

H.G., R.W., A.J., M.H., J.C.B., K.M.N., S.C., N.G.-Z., N.G. and C.T.: conceptualization. H.G., A.J., J.C.B. performed all experiments and investigations, H.G., R.W., A.J., M.H., J.C.B.: contributed to data curation and analysis, H.G., R.W., A.J., M.H., J.C.B., K.M.N., S.C., N.G.-Z., N.G. and C.T.: data interpretation, H.G., R.W., A.J., M.H., J.C.B., K.M.N., S.C., N.G.-Z., N.G. and C.T.: resources, H.G. and C.T.: writing. N. G.-Z., N.G., C.T.: supervision, C.T.: project administration, N. G.-Z., C.T.: funding acquisition. All authors read and approved the paper prior to submission.

## Declaration of generative AI and AI-assisted technologies in the writing process

During the writing of this work, the authors used ChatGPT and Gemini3 to improve sentence readability. All authors reviewed the manuscript after using this tool, edited the content, and take full responsibility for the content in this article.

## METHODS

### KEY RESOURCE TABLE

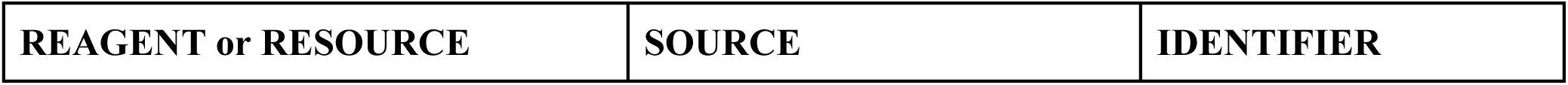
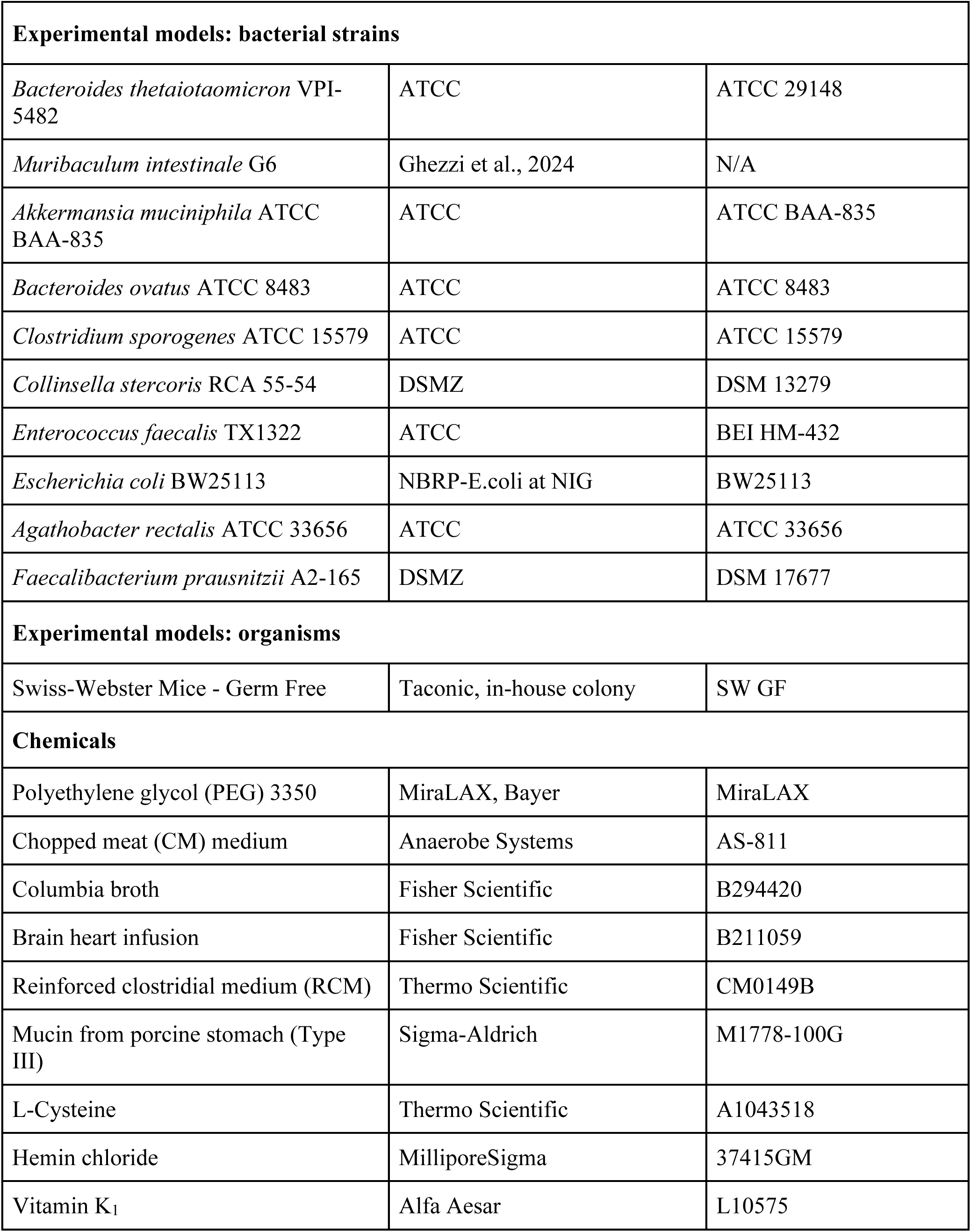
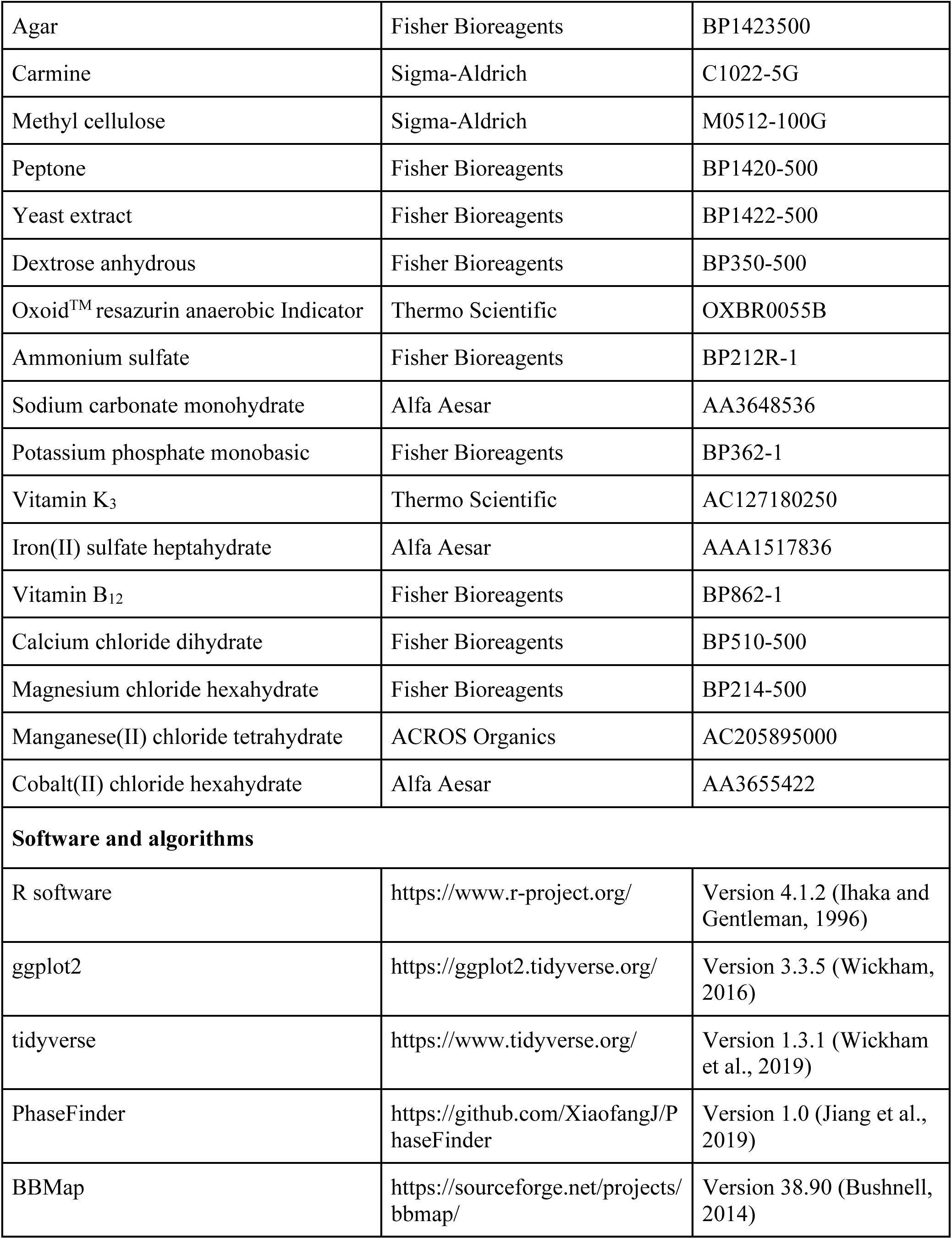
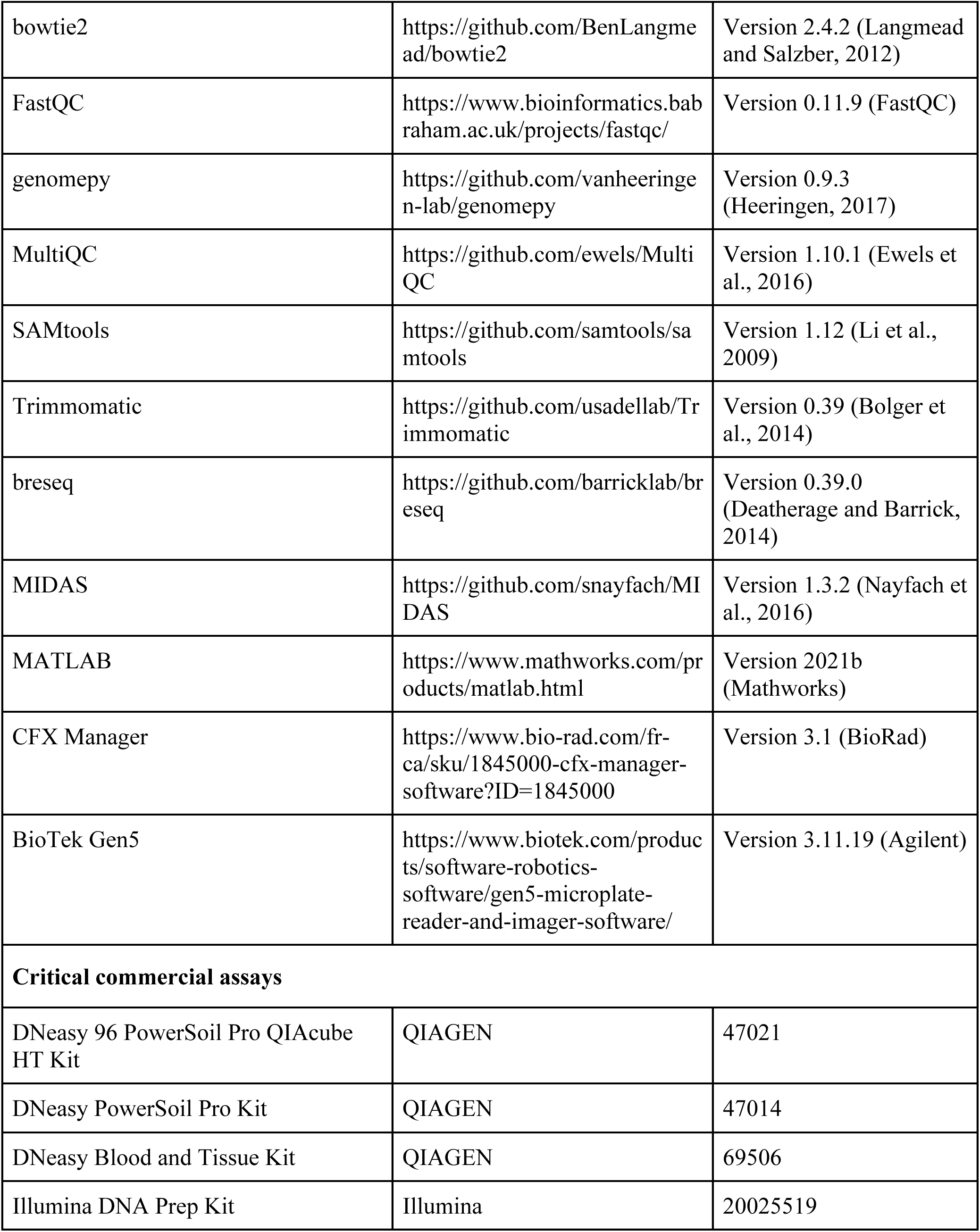
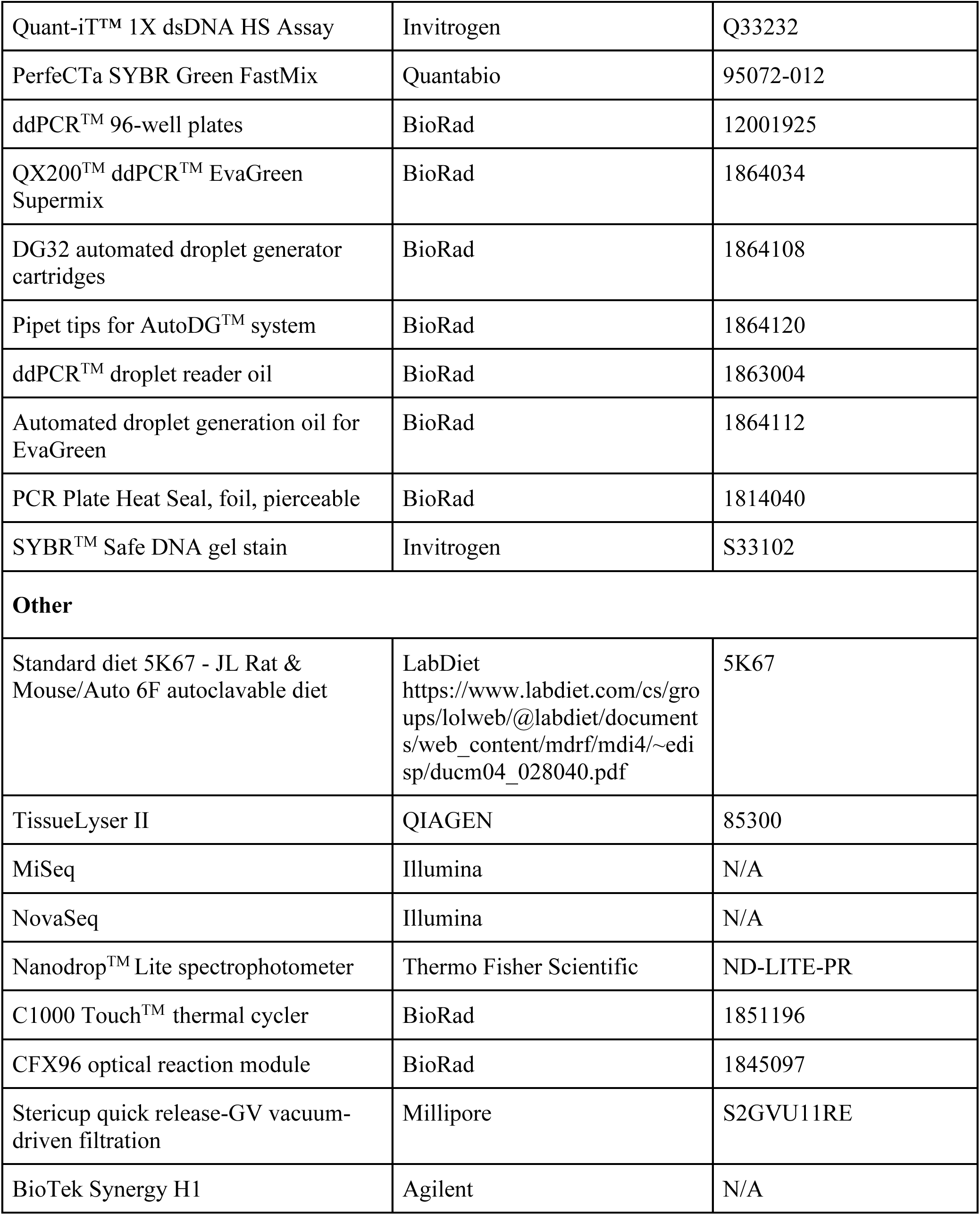

### EXPERIMENTAL MODELS

#### Animals

All animal experiments were performed according to protocol number (A19-0078) approved by the Animal Care Committee at the University of British Columbia and in direct accordance with the guidelines of the Canadian Council of Animal Care (CCAC). Mice used in this study were bred and maintained in a biosafety level-2 barrier at the GnotoCore, Centre for Disease Modelling rodent facility of the University of British Columbia, Canada.

Germ-free Swiss-Webster mice were maintained in gnotobiotic isolators or Ehret cages on a 12-hour light cycle (lights on between 07:00 and 19:00) and fed an autoclaved standard diet (Purina LabDiet 5K67). Port sterilization for material transfer into gnotobiotic isolators was achieved by spraying peracetic acid with an atomizer. All mice used were 10 to 12 weeks of age and co-housed with at least 3 mice per cage. Both genders were used and 3 to 5 littermates of the same sex were randomly assigned to each experimental group/condition.

At experimental end timepoint, mice were euthanised by carbon dioxide asphyxiation (primary), followed by cervical dislocation as a secondary method of euthanasia, prior to sample collection.

#### Bacterial strains and culture conditions

All bacterial strains used in this study were listed in the Key Resources Table and grown anaerobically (5% H_2_, 5% CO_2_, 90% N_2_) in a Coy anaerobic chamber at 37°C.

All bacterial strains, except *Akkermansia muciniphila* and *Faecalibacterium prausnitzii*, were cultured for ∼24-48h in supplemented Brain Heart Infusion (BHIS) plates (per liter: 37 g Bacto^TM^ Brain Heart Infusion and 15 g agar; media was autoclaved, cooled to 55°C and then supplemented with filter sterilised [0.20 μm] supplements: 1.5 mL of 333 mg mL^-1^ L-cysteine hydrochloride [5 g L-cysteine dissolved in 15 mL 1N HCl], 2 mL of 5 mg mL^-1^ hemin [50mg hemin dissolved in 1 mL 1N NaOH and 10mL ddH_2_O] and 1 mL of 1 mg mL^-1^ Vitamin K_3_ [100 μL Vitamin K_3_ in 10 mL 95% ethanol]). *A. muciniphila* was cultured for 6 days in BHIS + 1% mucin (w/v) plates (per liter: 37 g Bacto^TM^ Brain Heart Infusion, 15 g agar and 10 g mucin); media was autoclaved, cooled to 55°C and then supplemented as described above). *F. prausnitzii* was cultured for 2 days in supplemented Reinforced Clostridial Medium (RCM) plates (per liter: 37 g Thermo Scientific RCM [Dehydrated] and 15 g agar; media was autoclaved, cooled to 55°C and then supplemented as described above).

For inoculation in mice, single colonies of all bacterial strains, excluding *Collinsella stercoris, A. muciniphila* and *F. prausnitzii*, were individually inoculated in chopped meat (CM) media and grown overnight. *C. stercoris* single colonies were inoculated into supplemented Columbia (Columbia-S) growth media for approximately 24h (per liter: 35 g Difco^TM^ Columbia Broth; media was autoclaved, cooled to 55°C and then supplemented as described above). *A. muciniphila* single colonies were inoculated into BHIS + 1% mucin growth medium for approximately 24h (per liter: 37 g Bacto^TM^ Brain Heart Infusion and 10g mucin); media was autoclaved, cooled to 55°C and supplemented as described above. Media was centrifuged at 21130 rcf for 2 minutes, the supernatant was collected and filtered sterilised [0.20 μm]). *F. prausnitzii* single colonies were inoculated into supplemented RCM growth media for approximately 24h. After approximately 24 hours of growth, each isolated bacterial culture in stationary phase was pooled in a 1:1 ratio by volume and used to colonise germ-free mice by oral gavage.

#### Germ-free mouse inoculation

Germ free mice were colonised by oral gavage with 200 μl of 1:1 volume mix of isolated bacterial cultures (see above).

In the 10-member *in vivo* experiment, germ-free mice were colonised with the 10 microbes defined in the ‘GUT’ community by Ghezzi et al. (2024)^27^ while in the transit time experiment GF mice were bi-colonised with *B. thetaiotaomicron* and *M. intestinale* G6. Due to sparse engraftment of *A. rectalis* in the 10-member experiment, this microbe was excluded from *in vitro* assays. Following 6 and 4 weeks of equilibration for the 10-member community and bi-colonization experiments, respectively, each experimental group was administered autoclaved, filtered-sterilised, reverse-osmosis water supplemented with one the following polyethylene glycol (PEG) 3350 (Miralax) concentrations (w/v): 0%, 2.5%, 5%, 10% or 15%. PEG-supplemented water was administered for 6 consecutive days.

### *IN VIVO* SAMPLE PROCESSING AND ANALYSIS

#### PEG-supplemented water

PEG-supplemented water was made by supplementing standard cage drinking water (autoclaved reverse-osmosis water) with 0%, 2.5%, 5%, 10% or 15% (w/v) PEG 3350. PEG-supplemented water was thoroughly mixed on a stir plate for ∼5 minutes and then filter-sterilised (0.22 μm) using Stericup Quick Release-GV Vacuum-driven Filtration Systems (Millipore). Filter-sterilised water bottles were brought into the gnotobiotic isolators at least 2 days prior to day -1.

Immediately after transfer, the inner side of the bottlenecks was wiped with wet napkins to avoid contamination of water with peracetic acid that might have infiltrated during the port transfer into the isolator.

#### Samples and tissue collection

Fecal pellets were collected daily from each mouse for two days prior to administration of PEG-supplemented water (Day -1 and Day 0) and for the six days of treatment (Day 1 to Day 6). After dissection, cecal contents were collected and kept on ice until freezing at -80 °C for long-term storage.

#### Osmolality measurements

Osmolality measurements were performed on cecal contents immediately after sacrificing mice. Cecal osmolality was measured in duplicates using a Single-Sample Micro-Osmometer (Advanced Instruments OSMO1) and the averages reported.

#### Quantification of live fecal and cecal microbial abundance

Fecal and cecal samples were collected and immediately transported into an anaerobic chamber for quantification of colony forming units (CFUs) by spot plating. Duplicate sampling of 1 μL loops were serially diluted into filter sterilised (0.22 μm) PBS and spot plated on BHIS agar plates to measure total CFUs. Colony formation was monitored daily and CFUs were calculated by averaging the technical duplicates after ∼48 hours of anaerobic growth at 37 °C.

#### DNA extraction

Fecal samples collected daily throughout the experimental timelines were frozen at -80 °C until processed. High-throughput (HT) DNA extraction was performed simultaneously on all fecal and cecal samples. Samples were homogenised using a TissueLyser II (QIAGEN) for 10 minutes at 20 Hz and then processed with the DNeasy® 96 PowerSoil® Pro QIAcube® HT Kit (QIAGEN) and 96-well garnet PowerBead DNA plates (QIAGEN) according to the manufacturer’s extraction protocol. Following extraction, cecal DNA was quantified using a Quant-iT™ 1X dsDNA HS Assay Kit (Invitrogen) and Nanodrop. DNA integrity was confirmed on a 1% agarose gel.

#### Quantification of absolute microbial genomic copies

The absolute genomic abundance of individual GUT members in cecal samples was obtained with droplet digital PCR (ddPCR) using species-specific primers. Primer design and specificity checks were performed in Ghezzi et al. (2024)^27^. The oligonucleotide sequences can be found in Table S1 of this manuscript. All ddPCR reactions were run at Gut4Health (RRID:SCR_023673) as previously described^27^.

To quantify target copy numbers, cecal samples were first diluted in nuclease-free water (NF-H2O; Fisher Bioreagents) to a targeted range of ∼0–5,000 copies/µL. Droplet digital PCR assays were assembled in semi-skirted 96-well plates (Bio-Rad) and consisted of 11 µL of 2× QX200 ddPCR EvaGreen Supermix (Bio-Rad), 0.45 µL of forward/reverse primers (204 nM final concentration), 8.1 µL of NF-H2O, and 2 µL of diluted gDNA template. The reaction mixtures were partitioned into droplets via an Automated Droplet Generator (Bio-Rad) using DG32 cartridges, automated tips, and EvaGreen droplet generation oil (Bio-Rad). Plates were then enclosed with pierceable foil using a PX1 PCR Plate Sealer (Bio-Rad) and transferred to a C1000 Touch Thermal Cycler (Bio-Rad). The amplification protocol comprised an initial denaturation at 95 °C for 5 min, 40 cycles of denaturation at 95 °C for 30 s and annealing/extension at 62 °C for 1 min, followed by successive steps of 4 °C for 5 min and 90 °C for 5 min, with a final hold at 4 °C. A uniform ramp rate of 2 °C/s was applied across all steps. Droplet enumeration was conducted on a QX200 Droplet Reader (Bio-Rad) using specialized Droplet Reader Oil (Bio-Rad). Data visualization and manual amplitude thresholding were performed using QX Manager v2.0 software (Bio-Rad).

#### Library preparation and sequencing

Library construction was performed using Illumina DNA Prep by the Sequencing + Bioinformatics Consortium at the University of British Columbia, Vancouver, BC, Canada. Individual libraries were pooled together according to Illumina’s manufacturer protocol and sequenced on a NovaSeq500 2x150bp instrument.

#### Shotgun sequencing data analysis

Metagenomic reads were assessed for nucleotide quality using FastQC (v0.11.9) and the individual reports for each sample were aggregated using MultiQC (v1.10.1)^49^. Reads were processed with Trimmomatic (v0.39)^50^ to remove adapters (NexteraPE-PE.fa:2:30:10) and low-quality regions (LEADING:3 TRAILING:3 SLIDINGWINDOW:4:20 MINLEN:35). The mouse reference assembly (GRCm39/mm39) was downloaded with genomepy (v0.9.3)^51^ using -p UCSC -m hard and the processed reads were mapped against it with BBMap (v38.90)^52^ to remove sequences originating from the host. Microbial relative abundance and single nucleotide variants (SNVs) were obtained using MIDAS (v1.3.2) using default parameters^53^.

#### Whole gut transit time

Carmine red, a nonabsorbable dye, was used to measure gastrointestinal transit time in mice bi-colonised with *B. thetaiotaomicron* and *M. intestinale* G6. A carmine red solution was prepared by mixing 6% (w/v) carmine (Sigma-Aldrich) in 0.5% (w/v) methyl cellulose (Sigma-Aldrich) and heated nuclease-free water. This solution was autoclaved and imported into gnotobiotic isolators the day prior to the assay. Mice were acclimatized to the carmine red assay by gavaging empty feeding-needles followed by single-housing in empty cages for 3-4h. This acclimatization protocol was repeated twice in the week preceding transit measurements. Two days following the second acclimatization, mice were given 200 uL of the carmine red solution and immediately single-housed in empty cages. Fecal pellets were monitored for red color after 1h at 15 minutes intervals. Whole gut transit time was recorded as the time elapsed until the appearance of the first red pellet. Mice were sacrificed immediately after recording transit time.

### *IN VITRO* SAMPLE PROCESSING AND ANALYSIS

All *in vitro* assays performed to characterise bacterial physiological responses to osmotic stress were carried out anaerobically (5% H_2_, 5% CO_2_, 90% N_2_) at 37 °C. Osmotic stress was induced by supplementing the respective baseline media (see Methods above) with polyethylene glycol (PEG) 3350 (Miralax) to achieve the desired osmolality. Following supplementation, media was filter-sterilised and left in the anaerobic chamber for at least 12 hours before use.

#### Optical density measurements

For the characterization of *in vitro* growth under osmotic shock by optical density, single colonies of all bacterial strains, excluding *C. stercoris, M. intestinale, A.* muciniphila and *F.* prausnitzii, were inoculated in BHIS media and grown overnight. Liquid cultures were subcultured 1:100 and grown to exponential phase before being back diluted 1:100 into PEG-supplemented media for growth measurements. *C. stercoris* and *M. intestinale* colonies were inoculated into supplemented Columbia media and grown for ∼24h and ∼72h, respectively. *A. muciniphila* and *F. prausnitzii* colonies were inoculated into BHIS + 1% mucin and supplemented RCM, respectively, and grown for ∼24h. Liquid cultures were subcultured 1:10 and grown to exponential phase before being back diluted 1:100 into PEG supplemented media for growth measurements. Bacterial growth curves were collected using a BioTek™ Synergy™ H1 plate reader on the Biotek Gen5 v3.11.19 software at 37 °C inside an anaerobic chamber. Three to six replicates per bacterium were performed for each osmolality treatment. Optical density measurements at 600nm were collected every 5 minutes for at least 24h. The maximum growth rates r were obtained from the averaged curves using a MATLAB script^54^ which performs a least-squares fit of 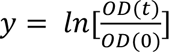 to the Gompertz equation^28^

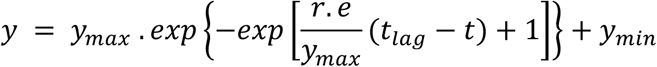

where *e* = *exp*(1), *y_min_* and *y_max_* are related to the maximum OD by *OD_max_* = *OD*(0). *exp*(*y_min_* + *y_max_*), and *t_la_*_g_ represents the lag time.

#### DNA extraction

All *in vitro* samples in this manuscript were processed for DNA extraction using the DNeasy Blood and Tissue kit (QIAGEN), following the manufacturer’s instructions. Samples were always eluted with 100 μL of elution buffer prior to use.

#### Passaging experiments

To investigate growth dynamics under osmotic stress in ancestral and evolved populations, *B. thetaiotaomicron* was streaked onto BHIS agar plates in triplicates and grown anaerobically at 37 °C for 24 hours. Single colonies were inoculated into BHIS liquid media, grown for 24 hours and then diluted 1:100 into BHIS media until exponential phase. These cells, referred to as the ancestral populations or “PRE”, were stored as glycerol stocks, processed for DNA extraction and qPCR, and further diluted 1:100 into either BHIS baseline media at ∼350 mOsm/Kg or BHIS supplemented with PEG at ∼950 mOsm/Kg for the passaging experiments. These samples were repeatedly cultured for 24 hours in either BHIS or BHIS-PEG and back-diluted 1:100 into respective fresh media for a total of 6 consecutive days. Samples were spot plated and optical density was quantified after 6, 12, and 24 hours of every passage event. After 6 days, liquid cultures from *B. thetaiotaomicron* populations were stored as glycerol stocks and processed for DNA extraction to perform shotgun sequencing and qPCR. BHIS evolved populations were referred to as “BHISe” and BHIS-PEG evolved ones as “PEGe”. PRE, BHISe, and PEGe *B. thetaiotaomicron* populations were double passaged in either BHIS or BHIS-PEG for a second 6-day event, as described above for the first passaging.

#### PBS mortality burden experiments

To quantify mortality burden, defined as the net decrease in viable colony-forming units, under osmotic shock *in vitro*, we inoculated a loopful of glycerol stocks from *B. thetaiotaomicron* ancestral and BHIS-evolved populations into BHIS liquid media (∼350 mOsm/Kg), and PEG-evolved populations into BHIS-PEG media (∼950 mOsm/Kg). Cultures in biological triplicates were grown anaerobically at 37 °C for 24 hours, then subcultured 1:100 into the respective media and grown until exponential phase. Cultures were then diluted 1:100 into PBS (∼300 mOsm/Kg) or PBS supplemented with PEG (∼950 mOsm/Kg). Live counts were obtained by spot plating 10-fold serial dilutions of each osmolality treatment on BHIS agar plates and monitoring colony forming units (CFUs) for 48 hours. Samples were spot plated immediately after dilution into PBS and PBS-PEG and then at regular intervals for varying amounts of time depending on the individual experiment. Viable colony-forming units at each timepoint were counted and normalised to the initial population baseline to calculate log10 mortality burden. BHIS agar plates were stored anaerobically at 37 °C for 48 hours before counting colonies and scraping the first row into 200 μL of PBS for DNA extraction and qPCR.

#### qPCR primer design and assays

Changes in proportion of *B. thetaiotaomicron* cells in the forward and reverse conformation of the CPS1 and PUL78/80 intergenic invertons were quantified using qPCR with the primers in Table S1. All qPCR reactions were performed in sealed, frosted 96-well PCR plate using a CFX96 optical reaction module on a C1000 TouchTM thermal cycler (BioRad) and analysed with CFX Manager v. 3.1 software (BioRad). Every qPCR reaction contained 12.5 μL of 2X PerfeCTa SYBR Green FastMix (QuantaBio), 0.5 μL of forward and reverse primers each (final concentration of 10 μM), 2 μL of template and 10.5 μL of nuclease-free water. qPCR amplification was achieved by running the following thermocycler program: 95 °C for 2m, 40 cycles of (95 °C for 10s, 62 °C for 30s, 72 °C for 30s) and 72 °C for 5m. Finally, a melting curve was generated by monitoring SYBR green fluorescence between 65 °C and 95 °C, with 0.5 °C increments and a hold of 5s. qPCR quantification was obtained using standard curves using extracted DNA from *B. thetaiotaomicron* pure cultures, which was quantified using Quant-iT™ 1X dsDNA HS Assay (Invitrogen), serially diluted 1:10 up to 10^6^ and run on all the qPCR reactions with the forward conformation primers of the respective variant, together with 2 non-template controls (NTCs).

#### Library preparation and sequencing

Library construction was performed using Illumina DNA Prep by Plasmidsaurus. Individual libraries were pooled together according to Illumina’s manufacturer protocol and sequenced on a 1.5B NovaSeq flow cell (2x150bp).

#### Shotgun sequencing data analysis

An average of 53,216,334 paired-end metagenomic reads per sample were quality controlled and trimmed as described above for the *in vivo* samples. Variant calling in *B. thetaiotaomicron* ancestral and evolved populations was performed with breseq (v.0.39.0)^55^. Briefly, trimmed reads from ancestral populations were used to apply fixed mutations from breseq clonal mode and reconstruct reference genomes starting from the *B. thetaiotaomicron* VPI-5482 genome (ASM1106v1). Single nucleotide polymorphisms in evolved and ancestral *B. thetaiotaomicron* populations were obtained using breseq in the polymorphism mode with a 1% frequency cutoff and a minimum of 10 supporting reads per strand. Additionally, *B. thetaiotaomicron* phase variants were identified using PhaseFinder (v.1.0)^30^. Briefly, the core ‘locate’, ‘create’, and ‘ratio’ functions of PhaseFinder.py were used with default parameters to identify changes in phase conformations in the breseq-updated reference genomes. Only invertons supported by more than 20 reads in the reverse conformation were retained for downstream analysis.

#### Transcriptional changes under osmotic stress

*B. thetaiotaomicron* transcriptional changes in the CPS1 and PUL78/80 loci were reported from publicly available datasets published in (McCallum, Burckhardt, et al., 2026)^35^.

##### B. thetaiotaomicron reporter activity

#### Genetic parts and plasmids

Plasmids were purified using a QIAprep Spin Miniprep Kit and sequence verified via whole-plasmid sequencing (Plasmidsaurus). Strains and plasmids can be found in Table S3. All primers and genetic parts used to generate these plasmids are listed in Table S4. Phusion® High-Fidelity DNA Polymerase was used according to manufacturer protocols for all parts constructed or amplified by PCR. PCR amplicons were purified in nuclease-free water using the QIAquick PCR Purification Kit prior to assembly. Golden Gate reactions were performed according to the protocol described below, and HiFi DNA assemblies were performed using the NEBuilder HiFi DNA Assembly (New England Biolabs) mix according to manufacturer protocols.

#### Golden Gate assemblies

All Golden Gate reactions were performed in 10 μL reactions containing: 0.33 μL BsaI HFv2, 0.33 μL T4 DNA Ligase, 1 μL 10x T4 DNA Ligase Buffer, 25 fmol Entry vector plasmid, 50 fmol each additional part, To volume with nuclease-free water.

Reactions were cycled as follows: 37 °C for 10 min 25 cycles of (37°C for 1.5 min, 16°C for 3 min), 37°C for 5 min, 80°C for 10 min, 12°C hold.

#### Competent cells and transformations

All cloning and plasmid construction was completed using either *E. coli* S17-1 λpir. For cloning into *E. coli* S17-1 λpir, cells were made chemically competent as described in (McCallum, Burckhardt, et al., 2026)^35^.

Briefly, to make TSS chemically competent cells, an overnight culture of *E. coli* S17-1 λpir was sub-cultured at a 1:100 ratio and grown until an OD_600_ of 0.4-0.5 (∼5 h). Cells were incubated on ice for 10 minutes before centrifuging at 3000 x *g* for 10 min at 4°C. Supernatant was removed and cells were resuspended in 10% of the original culture volume of TSS buffer (10% w/v PEG 3350; 0.03 M MgCl_2_; 5% v/v DMSO in LB-Miller broth). 200 μL aliquots of cells were stored at -80 °C until use. For transformation, 5 μL of assembly reaction was added to 100 μL thawed cells and incubated on ice for 30 min. Cells were then shocked at 42°C for 30 s and returned to ice for another 2 min. 900 uL of SOC recovery media was then added, and cells were incubated for 1 h at 37°C while shaking at 225 RPM. 300 μL of transformed cells were then plated on selective LB-agar plates and incubated at 37°C overnight.

#### P_TetR56_-based reporter plasmids for BT4299 upstream conformations

To generate constructs representing the two possible conformations of the BT4299 upstream invertible region, sequences flanking the IR where amplified via PCR from purified *B. thetaiotaomicron* gDNA. Approximately 1 kbp of sequence upstream and downstream of the IRs was amplified for each construct. Two custom synthetic gBlocks (Twist Bioscience) were designed, each containing one of the two possible orientations of the sequences between the IRs, and scrambled IR sequences to prevent inversions. The amplified flanking regions and gBlocks were introduced into a PCR-amplified pExchange backbone using HiFi assembly. Assembly HiFi reactions were transformed into TSS competent cells as described above.

The resulting pExchange constructs were subsequently used as templates to amplify 675 bp upstream from the BT4299 start codon. Primers were designed to introduce BsaI recognition sites and Golden Gate overhangs corresponding to the pEM576 entry vector (5’-CCGA-3’) at the 5’ end and *tetR* (5’-ACAT-3’) at the 3’ end. Amplified fragments were then cloned into the reporter plasmid pEM526 upstream of *tetR* using Golden Gate assembly as described above, generating reporter constructs containing each locked orientation of the BT4299 upstream region.

#### B. thetaiotaomicron conjugation

Wild type *B. thetaiotaomicron* was grown anaerobically overnight in BHIS broth at 37°C, then sub-cultured in 5 mL fresh BHIS at 1:50 dilution. *E. coli* S17-1 λpir containing assembled pNBU2 plasmids were grown overnight in LB-Amp, then sub-cultured in 5 mL fresh LB. Subcultures for both strains were grown ∼5 h, centrifuged at 3000 x *g* for 10 min and supernatant was discarded. *E. coli* and *B. thetaiotaomicron* pellets were serially resuspended in 1 mL of BHIS. 100 μL of resuspension was spot plated onto BHI+10% (v/v) horse blood agar plates and incubated at 37 °C aerobically for 14-16 h, followed by overnight anaerobic incubation at 37 °C. Conjugation spots were spread onto BHI+10% (v/v) horse blood agar plates with gentamicin (Gent; 200 μg/mL) and erythromycin (Erm; 25 μg/mL) and grown anaerobically for two nights. Single colonies were re-streaked onto BHIS+Erm+Gent plates and grown anaerobically for two nights. Single colonies were inoculated into liquid BHIS+Erm and grown overnight. For long-term storage, liquid cultures were mixed at a 1:1 ratio with 50% (v/v) glycerol and stored at -80°C.

#### Reporter i*n vitro* fluorescence measurements

Reporter constructs and no promoter control were streaked in BHIS + Erm plates and grown anaerobically overnight. Single colonies were picked and grown overnight in BHIS + Erm, then diluted 1:50 into either base BHIS media or media adapted to 700 mOsm/kg with PEG. Culture was then grown for ∼5 h until they reached an OD ∼0.4-0.7, removed from anaerobic incubation, harvested by centrifugation at 2500 x g for 10 minutes, and resuspended twice in PBS. After washing, cells were incubated at room temperature until 2 hours of aerobic exposure was reached. 200 μL of resuspended cells were transferred to a black, clear bottom 96-well plate and OD_600_, mGL (Ex/Em: 485/515 nm; Gain: 90) and mScarlet-I3 (Ex/Em: 570/600 nm; Gain 120) levels were measured using a Biotek Synergy H1 plate reader. mGL levels were normalized to the constitutive mScarlet-I3 levels, and activation for each construct was measured as the fold-decrease of the normalized GFP:RFP ratio relative to the no promoter control.

## SUPPLEMENTAL FIGURES

**FIGURE S1:**
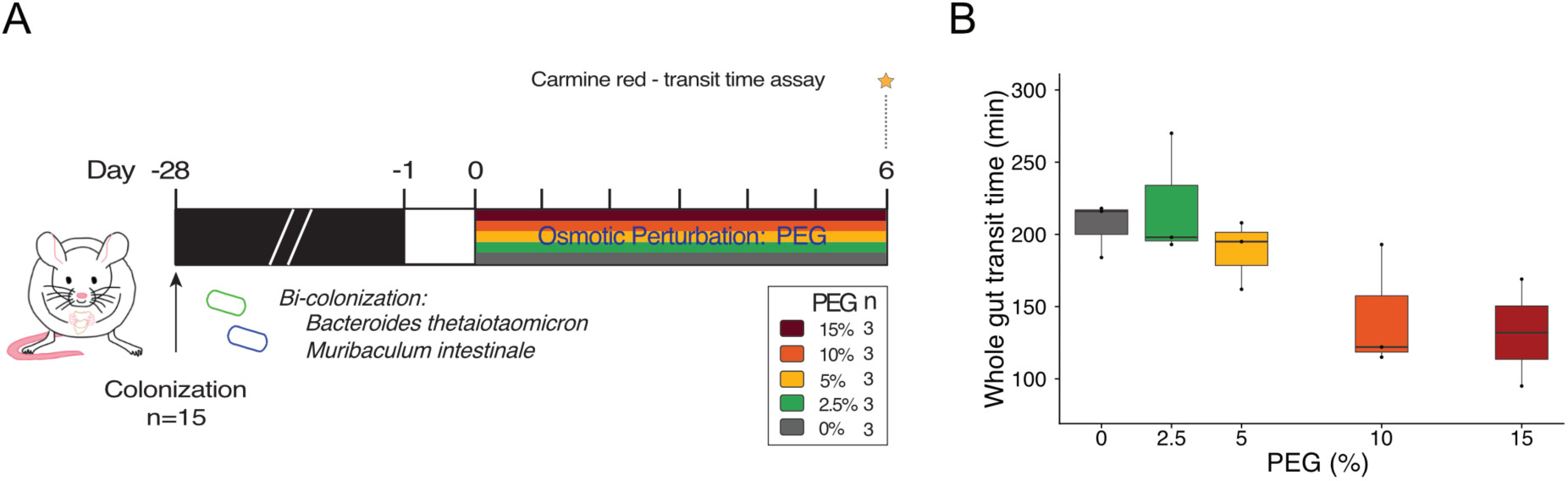
Whole gut transit time decreases following PEG treatment. **(A)** Experimental schematic and sample collection protocol. Germ-free mice (n=15) were colonised with *B. thetaiotaomicron* and *M. intestinale* and allowed to equilibrate for 28 days. Mice were treated with 0%, 2.5%, 5%, 10%, 15% PEG in drinking water (shown by color) *ad libitum* for 6 days after which they were gavaged with carmine red and monitored for dye in the stool to measure transit time. **(B)** Whole gut transit time measured after 6 days of treatments with 5 distinct PEG dosages n=3 each).

**FIGURE S2:**
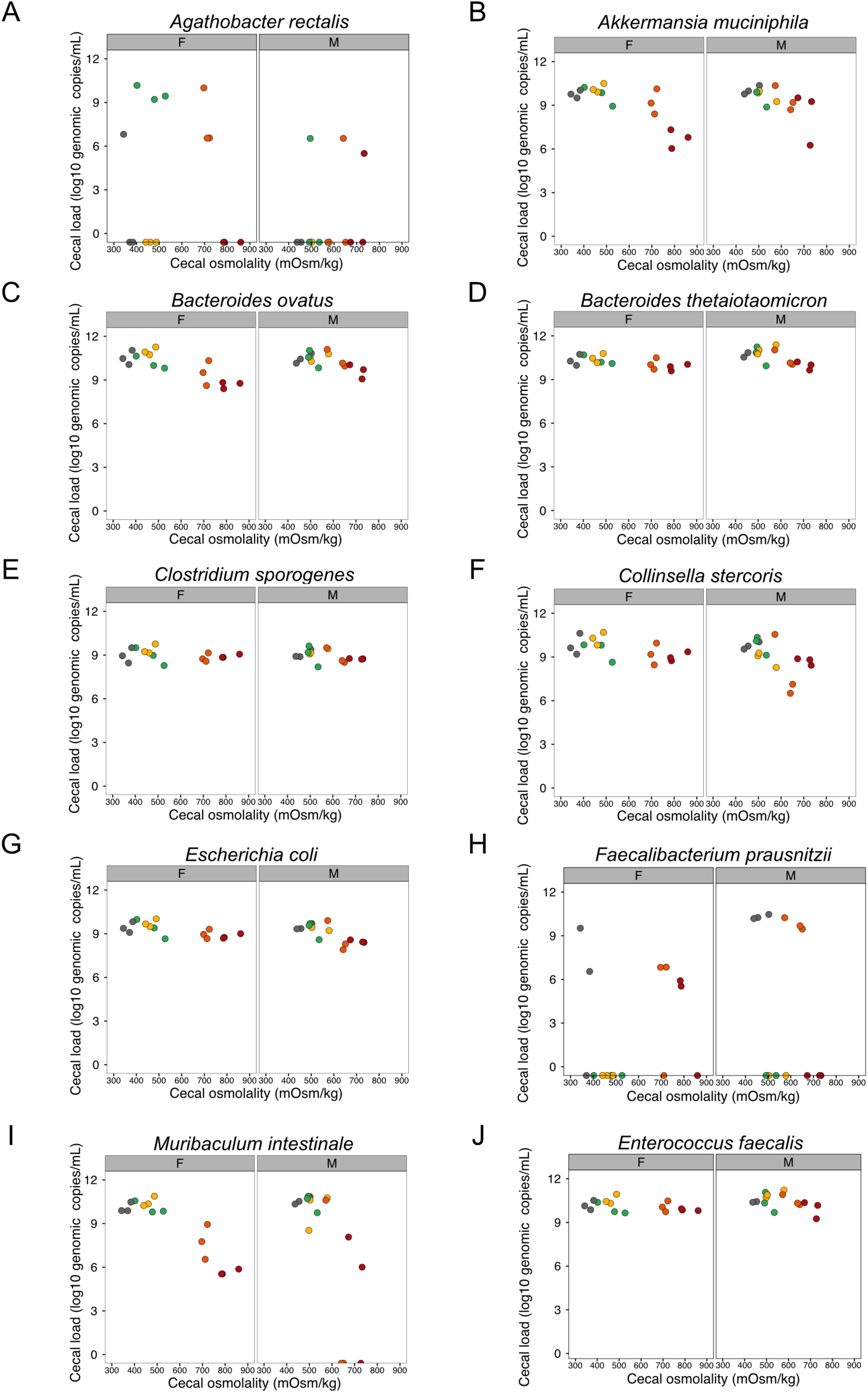
Species-specific absolute quantification reveals distinct osmotolerance levels in a defined 10-member community. Absolute abundance (log_10_ genomic copies/mL) measured by species-specific droplet digital PCR (ddPCR) from cecal samples collected after 6 days of PEG treatment (see Fig. 1A for experimental schematic). Individual panels display data for each community member: **(A)** *Agathobacter rectalis*, **(B)** *Akkermansia muciniphila*, **(C)** *Bacteroides ovatus*, **(D)** *Bacteroides thetaiotaomicron*, **(E)** *Clostridium sporogenes*, **(F)** *Collinsella stercoris*, **(G)** *Escherichia coli*, **(H)** *Faecalibacterium prausnitzii*, **(I)** *Muribaculum intestinale*, and **(J)** *Enterococcus faecalis*. Individual points represent individual mice plotted against their respective cecal osmolality (mOsm/kg). Plots are faceted by the sex of the mice (F, female; M, male). Points are color-coded by PEG dosage (0%, 2.5%, 5%, 10%, and 15% PEG) according to the color scheme in Fig. 1A.

**FIGURE S3:**
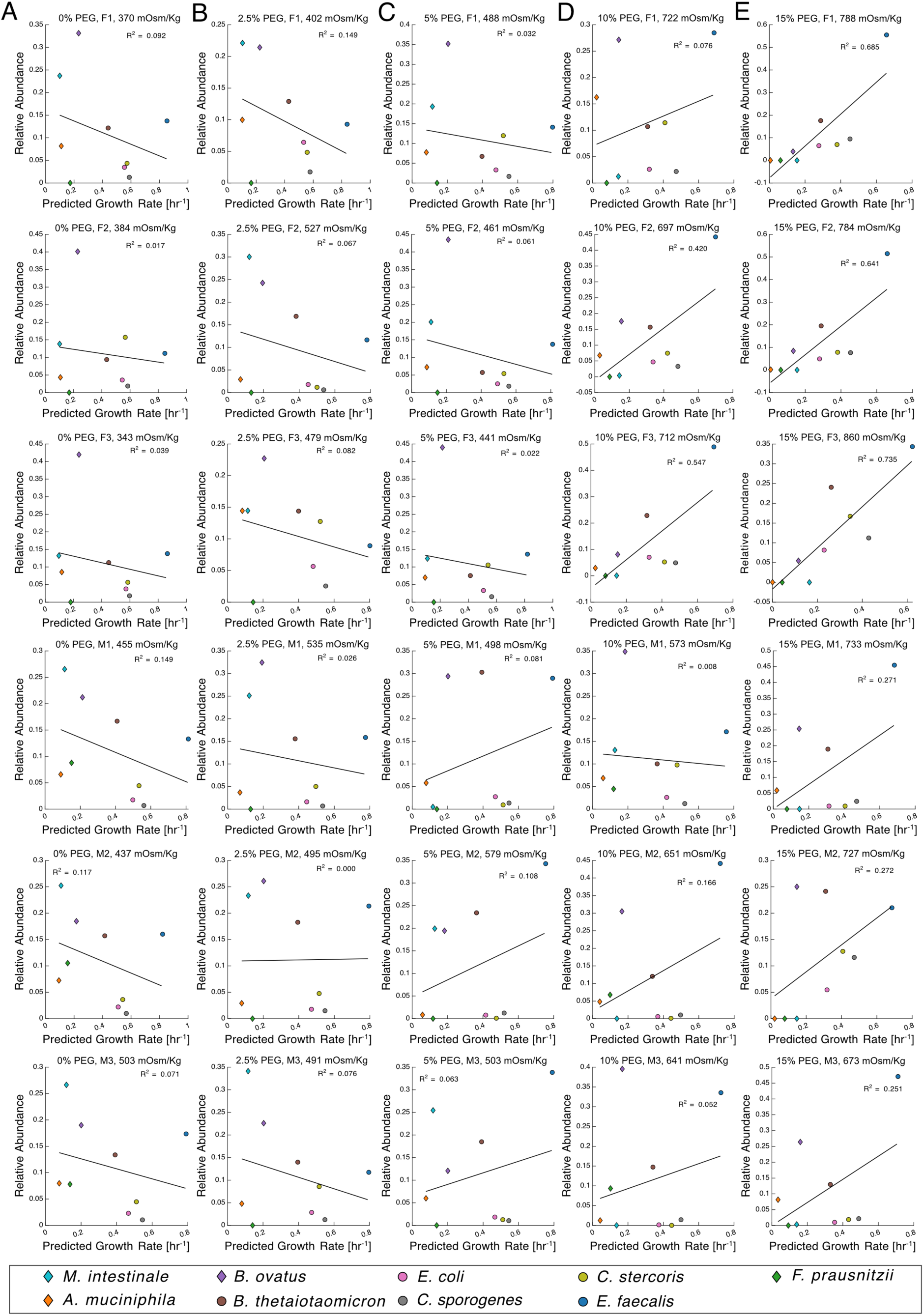
Mouse-specific linear regressions between *in vivo* relative abundance and predicted *in vitro* growth rate. Columns represent the experimental PEG treatment groups in the experiment outlined in Fig. 1A: (**A**) 0% PEG, (**B**) 2.5% PEG, (**C**) 5% PEG, (**D**) 10% PEG, and (**E**) 15% PEG. Rows represent individual mice, labeled by sex and numerical identifier (F1-F3 for females; M1-M3 for males) along with their measured cecal osmolality (mOsm/kg) at endpoint. Within each panel, the *in vivo* relative abundance of the 10 community members is plotted against their predicted *in vitro* growth rate at that mouse’s specific cecal osmolality. Individual points represent distinct bacterial strains, color-coded by species according to the legend at the bottom. The individual Pearson correlation coefficients derived from these 30 independent linear regressions were compiled to generate the summary correlation profile shown in Fig. 1G.

**FIGURE S4:**
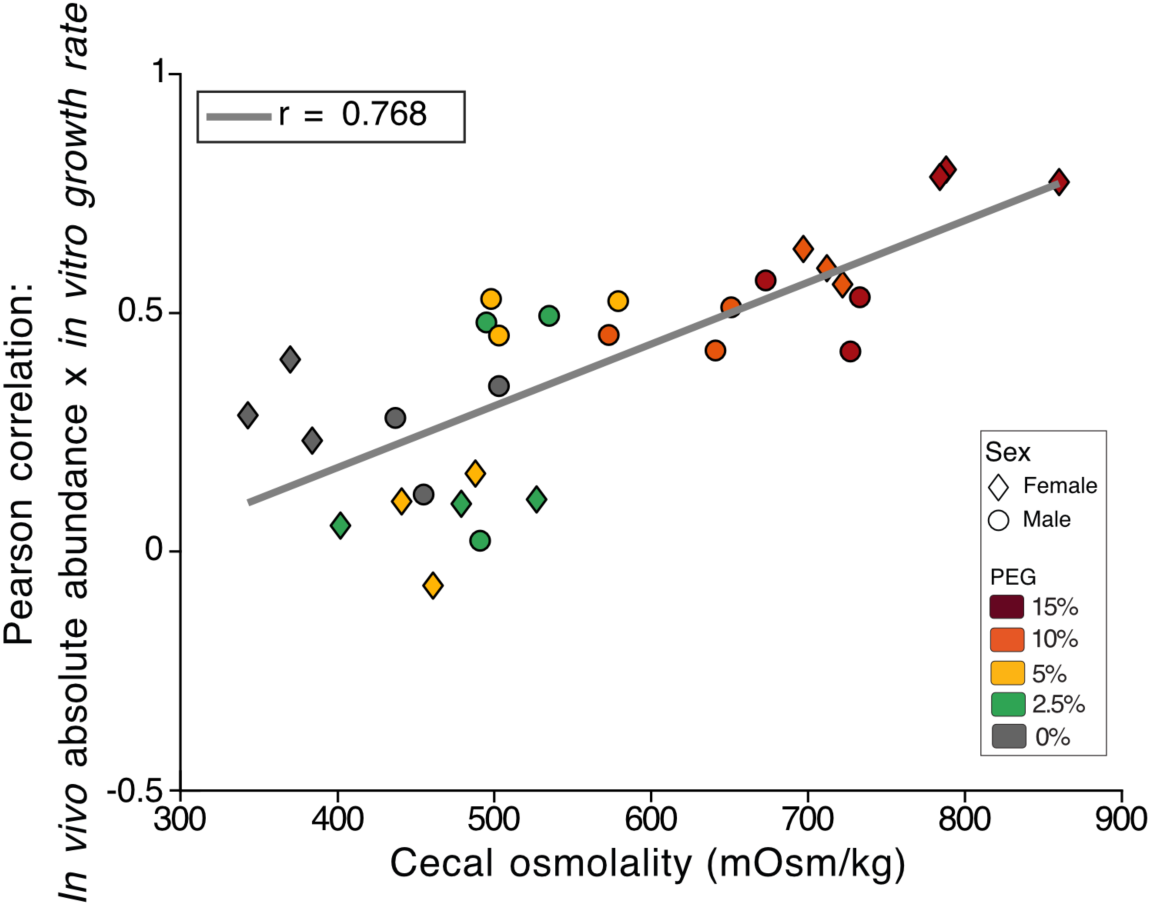
Pearson correlation between predicted *in vitro* growth rate and *in vivo* absolute abundance across cecal osmolalities. Pearson correlation coefficients calculated from linear regressions between species-specific *in vivo* absolute abundance measured via ddPCR (see Fig. S2) and predicted *in vitro* maximum growth rates. Individual points represent individual mice plotted against their measured cecal osmolality. Color denotes PEG dosage according to the scheme in Fig. 1A, and shape represents the sex of the mice (triangles for females, circles for males).

**FIGURE S5:**
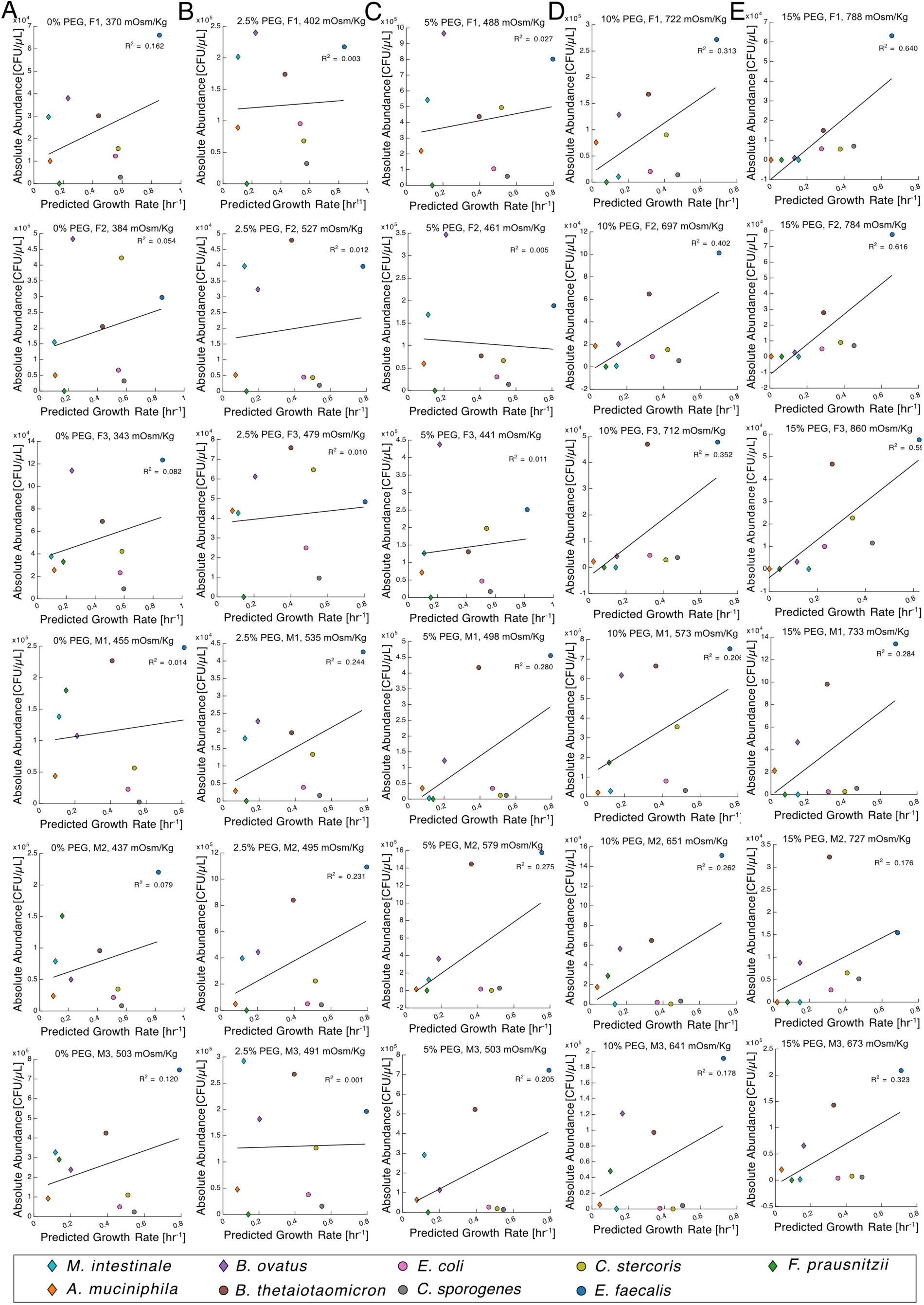
Mouse-specific linear regressions between *in vivo* absolute abundance and predicted *in vitro* growth rate. Columns represent the experimental PEG treatment groups in the experiment outlined in Fig. 1A: (**A**) 0% PEG, (**B**) 2.5% PEG, (**C**) 5% PEG, (**D**) 10% PEG, and (**E**) 15% PEG. Rows represent individual mice, labeled by sex and numerical identifier (F1-F3 for females; M1-M3 for males) along with their measured cecal osmolality (mOsm/kg) at endpoint. Within each panel, the *in vivo* absolute abundance of the 10 community members is plotted against their predicted *in vitro* growth rate (hr^-1^) at that mouse’s specific cecal osmolality. Individual points represent distinct bacterial strains, color-coded by species according to the legend at the bottom. The individual Pearson correlation coefficients derived from these 30 independent linear regressions were compiled to generate the summary correlation profile shown in Fig. S4.

**FIGURE S6:**
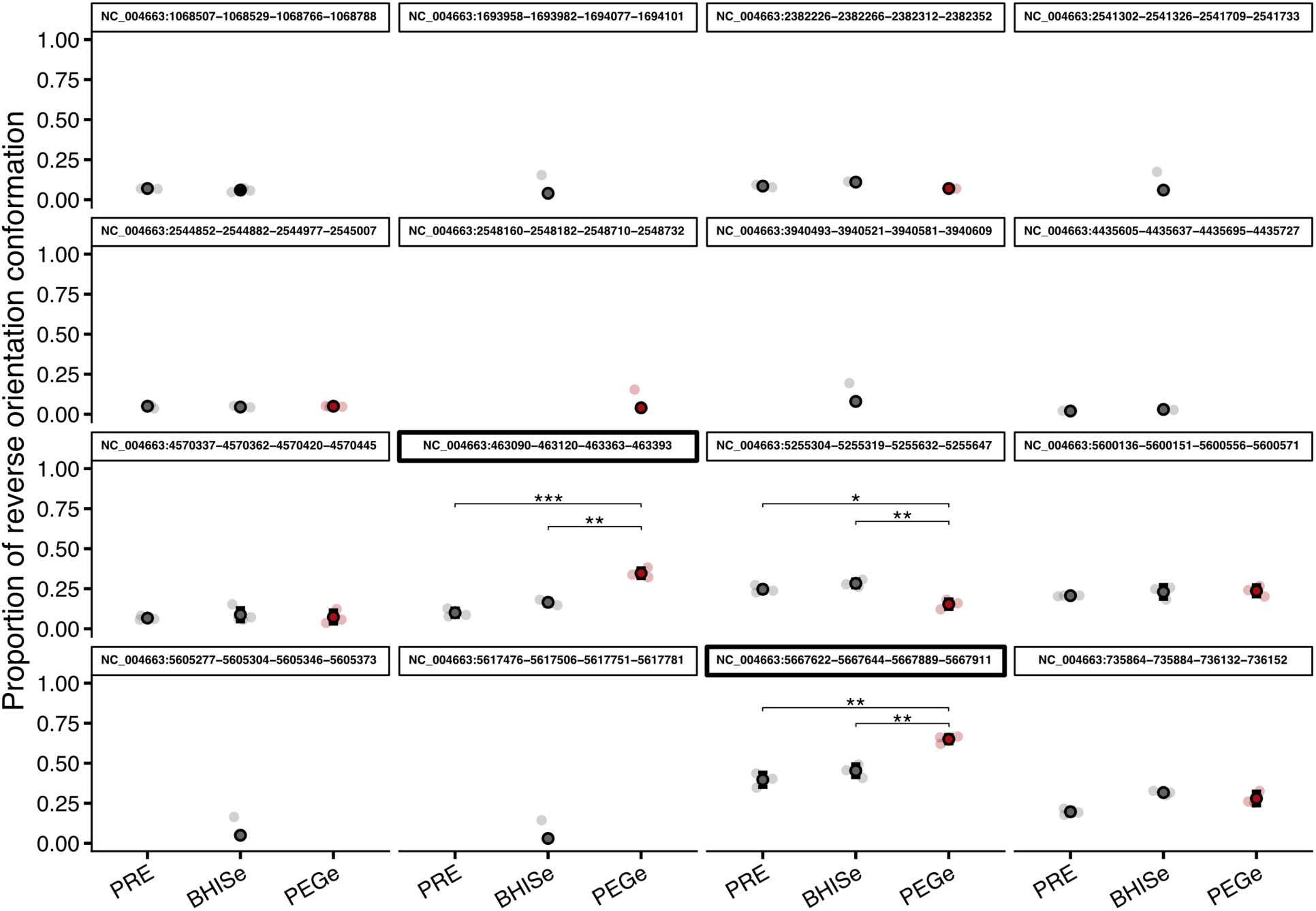
**PhaseFinder screening of invertible promoters across *B. theta****iotaomicron* **populations.** Proportion of reverse promoter conformations across all candidate phase variants identified via PhaseFinder after the first passaging round. Only invertons with a minimum of 20 reads mapping to the reverse conformation in at least one population are displayed. Faint points represent individual biological replicates (n=3), bold points indicate the mean, and error bars denote the standard deviation. Panels outlined with thick black borders highlight the only two invertons that exhibited a significant increase in reverse conformation frequency in PEGe populations compared to controls: near the CPS1 cluster (third row, second column) and the PUL78/80 operon (fourth row, third column). Statistical significance was determined by a two-tailed, paired t-test (*p < 0.05, **p < 0.01, ***p < 0.001).

## SUPPLEMENTAL TABLES

**TABLE S1:** Oligos used in this study.

**TABLE S2:** Raw breseq output files from shotgun sequencing of passaged *B. thetaiotaomicron* populations, related to Figure 2.

**TABLE S3:** Genetic parts and primers used in this study to engineer wild type *B. thetaiotaomicron*, related to Figure 3.

**TABLE S4:** Strains and plasmids used in this study to engineer wild type *B. thetaiotaomicron*, related to Figure 3.

